# LINE1 RNA dysregulation impairs chromatin accessibility in C9ORF72- and TDP-43-linked ALS/FTD

**DOI:** 10.1101/2025.09.30.679260

**Authors:** Yini Li, Xiaoyang Dou, Yu Xiao, Zhe Zhang, Yingzhi Ye, Noelle Wright, Koping Chang, Chang Liu, Juan C Troncoso, Chuan He, Shuying Sun

## Abstract

The long interspersed element-1 (LINE1) retrotransposon RNAs are abnormally elevated in various neurodegenerative disorders, but their pathogenic roles remain unclear. Here we investigated the mechanism of LINE1 RNA accumulation and its function in amyotrophic lateral sclerosis (ALS) and frontotemporal dementia (FTD) associated with C9ORF72 repeat expansion and TDP-43 loss-of-function, the leading causes of familial and sporadic forms of these neurodegenerative diseases. We show that LINE1 RNA is dysregulated due to an impaired nuclear exosome targeting (NEXT) degradation pathway. Its elevation epigenetically increases chromatin accessibility, enhancing global transcription via a retrotransposon-independent mechanism. Reducing LINE1 RNA mitigates chromosomal abnormalities and improves the survival of disease-relevant neurons. These findings uncover an essential noncoding RNA function and regulatory mechanism of LINE1 in neurons, providing insights into disease pathogenesis and highlighting potential therapeutic targets for neurodegenerative diseases.

## Main Text

The human genome contains millions of repetitive retrotransposon sequences, with long interspersed elements (LINEs) being the most abundant, represented by about 0.9-1 million copies^1^. Retrotransposons are normally repressed in somatic cells^2^. However, elevation of retrotransposon expression, especially LINE1, has been associated with aging and many neurodegenerative diseases, including amyotrophic lateral sclerosis (ALS), frontotemporal dementia (FTD), and Alzheimer’s disease^3–7^. Retrotransposition-competent full-length LINE1 contains a 5’ bidirectional promoter, two open reading frames (ORFs) that encode RNA chaperone (ORF1p) and endonuclease/reverse transcriptase (ORF2p), and a 3’ untranslated region with a weak polyadenylation signal^8,9^. Although LINE1 is the only known active autonomous transposable elements in the human genome, a large majority of LINE1 elements have undergone truncations and mutations, resulting in loss of retrotransposition activity^10^. Despite the one active LINE1 subfamily, 99.9% of LINE1s are retrotransposition-incompetent. Increasing studies have shown that LINE1 element in the transcribed RNA (hereafter we collectively refer to as LINE1 RNA) plays a significant role in regulating epigenetic identity and chromatin structure in mouse embryonic stem cells (mESCs), independently of retrotransposition^11,12^. Although elevation of LINE1 RNAs are widely reported in neurogenerative diseases, the possible non-coding function of LINE1 RNA is largely neglected. Moreover, the mechanism underlying the abnormal LINE1 RNA elevation is yet to be resolved.

*N*^6^-methyladenosine (m^6^A) is the most prevalent internal RNA modification in eukaryotes. m^6^A is installed by methyltransferases METTL3 and METTL14, removed by demethylases FTO and ALKBH5, and recognized by an extended list of RNA-binding proteins (RBPs) referred to as “readers”^13^. m^6^A modifies both coding and non-coding RNAs^14^. Most recently, LINE1 RNAs have been found to be heavily modified and regulated by m^6^A in mESCs and early embryos^15–17^. We previously discovered that m^6^A is globally reduced in C9ORF72-ALS/FTD^18^, the most common familial form of ALS and FTD, caused by a GGGGCC hexanucleotide repeat expansion in the human *C9ORF72* gene^19^. Notably, RNA-seq data from studies on C9ORF72-ALS/FTD patient revealed significant LINE1 elevation in patients’ brain tissues^3^. These data raise the intriguing possibility that global m^6^A reduction could lead to LINE1 dysregulation, potentially contributing to disease progression in C9ORF72-ALS/FTD.

TDP-43, encoded by the TAR DNA-binding protein (*TARDBP*) gene, is an RNA-binding protein (RBP) with multiple functions in RNA processing^20^. Nuclear clearance and cytoplasmic aggregation of TDP-43 is a major pathological hallmark of sporadic ALS and FTD, accounting for 97% of all ALS cases and 45% of all FTD cases^21,22^. Additionally, TDP-43 proteinopathy occurs in about 20-60% Alzheimer’s^23,24^. It is increasingly suggested that the loss of nuclear TDP-43 function is likely an early pathological event driving disease progression^25,26^. TDP-43 has been shown to bind to LINE1 RNAs^27^, which contains the enriched binding motif UGUGU^28^. Chronic deficiency of TDP-43 is implied in LINE1 RNA-associated R-loop formation^29^. Loss of nuclear TDP-43 is associated with increased LINE1 in patients’ tissues^30^, though the mechanisms are not yet understood.

The elevation of mobile LINE1 and retrotransposition activity has been well documented in cancer cells^31^, but the regulation and functional role of LINE1 in neurons and neurodegenerative diseases remain unclear. Here we show that LINE1 RNA is predominantly associated with chromatin in neurons and poorly encodes the LINE1 proteins required for the retrotransposition activity. Its expression becomes dysregulated due to impaired RNA degradation via the NEXT pathway, independently triggered by either TDP-43 reduction or C9ORF72 repeat expansion — a convergent pathogenic mechanism implicated in the two forms of ALS/FTD. Elevated LINE1 RNA epigenetically increases chromatin accessibility and enhances global transcription. Reducing LINE1 RNA alleviates chromatin abnormalities and improves the survival of patient-derived and disease-relevant neurons. These findings uncover a previously unrecognized regulatory RNA function of LINE1 in neurons, providing new insight into disease mechanisms and highlighting potential therapeutic targets for neurodegenerative diseases.

## Results

### LINE1 RNA is chromatin-enriched and poorly encodes proteins in iPSNs

To investigate the possible function of LINE1 RNA in neurons, we examined its sub-cellular localization in human-derived induced pluripotent stem cell (iPSC)-differentiated neurons (iPSNs). We fractionated the iPSNs into cytoplasmic, nucleoplasmic, and chromatin fractions (Extended Data Fig. 1a) and quantified total LINE1 RNA by RT-qPCR using primers targeting a common region of LINE1 RNAs^32–34^. RNAs were treated with DNase to remove genomic DNA, and control reactions without reverse transcriptase showed negligible signal. The LINE1 RNA levels in each subcellular compartment were normalized to spike-ins. LINE1 RNA was highly enriched in the chromatin-associated fraction (Fig. 1a), indicating a potential role in chromatin-mediated regulation.

**Fig.1:**
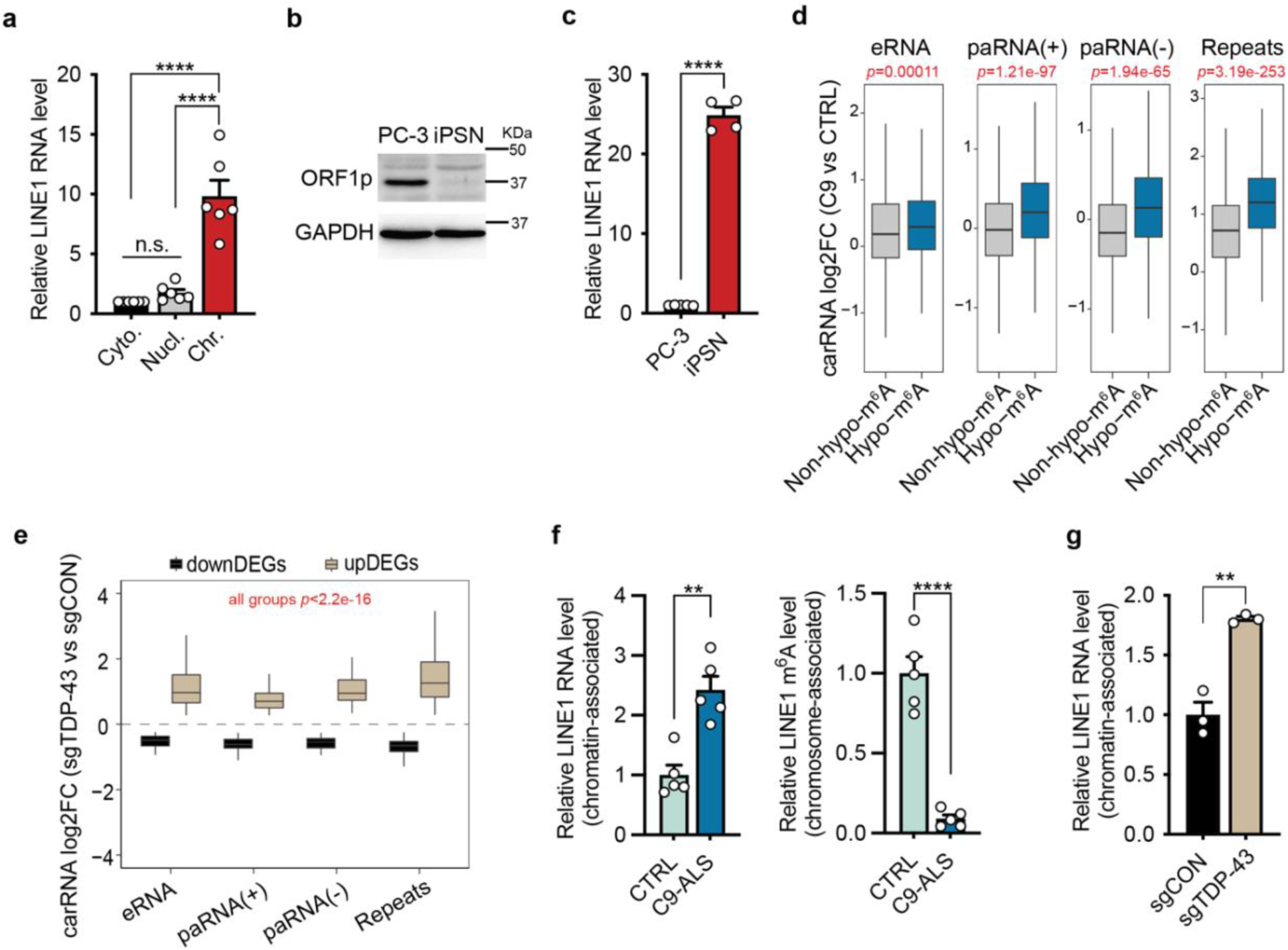
LINE1 RNA is chromatin-enriched and increased in C9ORF72-ALS/FTD and TDP-43-reduced neurons. **a,** Subcellular distribution of LINE1 RNA in control iPSNs, measured by RT-qPCR. The relative expression of LINE1 RNA was normalized to the spike-in supplemented in each fraction. Cyto., cytoplasmic, Nucl., nucleoplasmic, Chr., chromatin-associated fractions. Points represent individual control iPSN lines (n=6). P-values were calculated by one-way analysis of variance (ANOVA) with Tukey’s multiple comparisons. **b**, Western blotting of ORF1p in PC-3 cancer cells and iPSNs. GAPDH was blotted as internal control. **c**, Relative LINE1 RNA expression level in PC-3 and iPSNs, measured by RT-qPCR. Points represent different batches of PC-3 culture (n=5) and individual control iPSN lines (n=4). P-values were calculated by two-tailed *t* test. **d**, Boxplots showing carRNA expression changes with or without hypo-m^6^A in C9ORF72-ALS/FTD patient iPSNs. eRNA: enhancer RNA; paRNA(+): sense-strand promoter-associated RNA; paRNA(-): antisense-stranded promoter-associated RNA; Repeats: repeat RNA. n=4 control lines and n=4 C9 lines. P-values were calculated by two-tailed Student’s *t* test. Box plots indicate the interquartile range with the central line representing the median, and the vertical lines extend to the extreme values in the group. **e**, Boxplots showing carRNA expression changes in TDP-43-depleted i^3^Neurons. n=3 control replicates (sgCON) and n=3 TDP-43-knockdown replicates (sgTDP-43). The differential expression gene cutoff was P_adj_<0.05. **f**, Relative expression levels (left) and m^6^A levels (right) of LINE1 RNA, measured by RT-qPCR and MeRIP-RT-qPCR in iPSNs of controls and C9ORF72-ALS/FTD patients. Points represent individual lines (*n*=5 lines per group). **g**, Relative LINE1 RNA expression levels measured by RT-qPCR in control and TDP-43-reduced i^3^Neurons (n=3 biological replicates per group). P-values were calculated by two-tailed *t* test. Data are mean ± s.e.m. **p < 0.01, ****p < 0.0001.

We next measured the total LINE1 RNA and protein in neurons and compared them to the PC-3 prostate cancer cell line (Fig. 1b,c), which exhibits high LINE1 protein expression and retrotransposition activity^35^. As expected, there was robust expression of LINE1 ORF1p in PC-3 cells (Fig. 1b). Strikingly, the RNA level of LINE1 in iPSNs was drastically higher than that in PC-3 cells (Fig. 1c), yet the ORF1p protein was barely detectable (Fig. 1b). This finding indicates that, different from cancer cell, the LINE1 RNA does not effectively encode proteins in neurons. It is known that LINE1 proteins has a strong *cis-*preference on binding to the RNA that they are translated from to execute retrotransposition^36^. The relatively low protein levels despite abundant RNA suggest minimal retrotransposition activity. Combined with the enrichment of LINE1 RNA in chromatin fractions (Fig. 1a), these results suggest the large proportion of the untranslated LINE1 may serve as a regulatory RNA in neurons, rather than carrying out its canonical retrotransposition function.

### LINE1 RNA is elevated in both C9ORF72-ALS/FTD and TDP-43-reduced neurons

We next examined whether LINE1 RNAs are altered in the chromatin fractions of iPSNs derived from C9ORF72-ALS/FTD patients or neurons with TDP-43 reduction. We examined the chromatin-associated regulatory RNAs (carRNAs) that include promoter-associated RNA, enhancer RNA, and RNA transcribed from transposable elements (repeat RNA)^16^. For C9ORF72-ALS/FTD, we isolated RNA from the chromatin fractions of multiple lines of iPSNs from both non-neurological controls and patients. As the global m^6^A on mRNAs was previously found to be reduced in C9ORF72-ALS/FTD iPSNs^18^, we performed m^6^A-immunoprecipitation sequencing (MeRIP-seq) to assess both m^6^A modification levels and total levels of carRNAs. m^6^A was reduced in all the three types of carRNAs in C9ORF72-ALS/FTD iPSNs (Extended Data Fig. 1b). Compared to non-hypomethylated carRNAs (non-hypo-m^6^A), m^6^A hypomethylated carRNAs (hypo-m^6^A) showed significantly higher expression in C9ORF72-ALS/FTD, most profoundly in the LINE1-containing repeat RNA group (Fig. 1d). Moreover, the majority of carRNAs were upregulated, with LINE1 RNA showing the most significant increase (Extended Data Fig. 1c). These results indicate that chromatin-associated LINE1 RNA is elevated in C9ORF72-ALS/FTD, and the increase is associated with m^6^A hypomethylation.

To study the chromatin-associated LINE1 RNA changes caused by TDP-43 loss of function, reflecting sporadic neurodegeneration conditions, we used i^3^Neurons^37^ and suppressed TDP-43 expression by sgRNA-mediated gene silencing (Extended Data Fig. 1d). We performed total RNA sequencing on the carRNAs and compared the changes between TDP-43 knockdown neurons and negative controls. We found that TDP-43 reduction led to significant expression changes of carRNAs (Fig. 1e), including a marked upregulation of LINE1 RNAs (Extended Data Fig. 1e), similar to the changes observed in C9ORF72-ALS/FTD iPSNs (Extended Data Fig. 1c).

We verified the LINE1 RNA changes in C9ORF72-ALS/FTD and TDP-43 knockdown neurons by RT-qPCR (Fig. 1f,g). We also confirmed the LINE1 RNA elevation in postmortem motor cortex tissues from C9ORF72-ALS/FTD patients (Extended Data Fig. 2a), accompanied by a reduced m^6^A level (Extended Data Fig. 2b). Interestingly, most of the upregulated LINE1 elements in neurons under both conditions originate from evolutionarily ancient subfamilies^38^ (Extended Data Fig. 2c,d). As only the young full-length LINE1 maintains retrotransposition activity^38^, this result supports a non-canonical function of LINE1 RNA that is uncoupled from LINE1 retrotransposition function.

### Abnormal LINE1 RNA accumulation results from reduced NEXT-mediated RNA decay

To examine whether increased LINE1 RNA is associated with a prolonged RNA decay rate, we measured the half-life of LINE1 RNA in both disease conditions. We treated the cells with actinomycin D to halt de novo transcription, collected RNA samples at 0 hours, 3 hours, and 6 hours post-treatment, and performed LINE1 RT-qPCR. LINE1 RNA showed prolonged half-life in both C9ORF72-ALS/FTD iPSNs and TDP-43 knockdown i^3^Neurons (Fig. 2a,b). Nuclear exosome targeting (NEXT) complex is a nuclear RNA decay machinery that has been shown to promote LINE1 RNA degradation in mESCs^16^. Therefore, we investigated its role in regulating LINE1 RNA in neurons. RNA-immunoprecipitation with the NEXT core component, ZCCHC8, showed that ZCCHC8 strongly binds to LINE1 RNA in neurons (Fig. 2c). Additionally, knockdown of ZCCHC8 in neurons led to significant increase of LINE1 RNA (Fig. 2d), confirming that NEXT degrades LINE1 RNA under normal conditions, preventing its accumulation.

**Fig.2:**
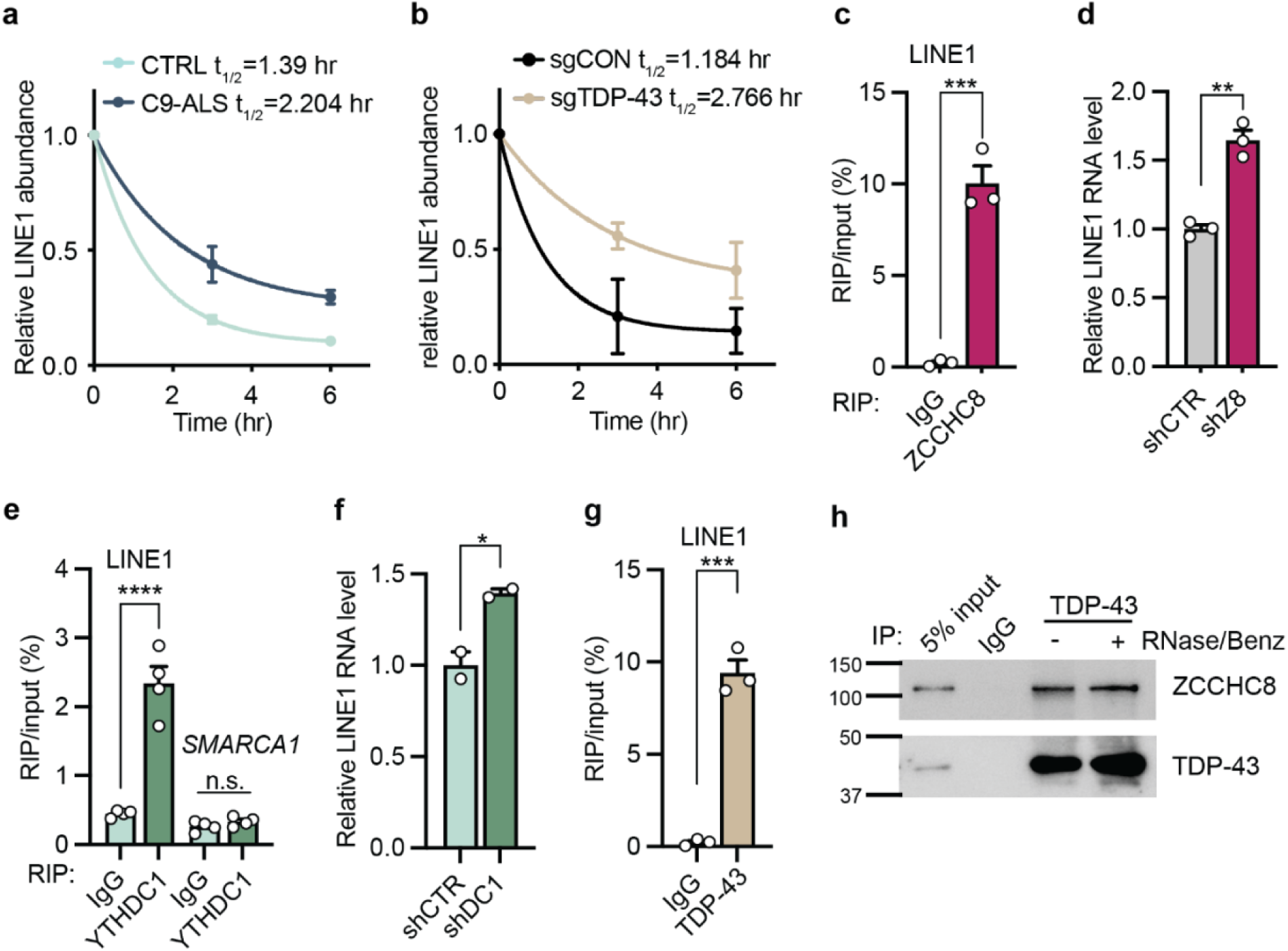
Defective NEXT-mediated LINE1 RNA decay in C9ORF72-ALS/FTD and TDP-43-reduced neurons. **a-b**, Half-life of the LINE1 RNA in (a) control and C9ORF72-ALS/FTD iPSNs, and (b) sgCON and sgTDP-43 i^3^Neurons. The RNA level was measured by RT-qPCR at each time point and normalized to the data at time 0. *n*=3 individual control or C9ORF72-ALS/FTD iPSN lines. *n*=5 biological replicates of sgCON or sgTDP-43. **c**, ZCCHC8-associated LINE1 RNA measured by RIP-RT-qPCR. Points represent individual iPSN lines. P-values were calculated by two-tailed *t* test. **d**, Relative LINE1 RNA expression measured by RT-qPCR upon knockdown of ZCCHC8 in i^3^Neurons. Points represent 3 biological replicates. P-values were calculated by two-tailed Student’s *t* test. **e**, YTHDC1-RIP with RT-qPCR of LINE1. Non-m^6^A modified *SMARCA1* transcript was used as negative control. Points represent individual iPSN lines. P-values were calculated by two-tailed Students’s *t* test. **f**, Relative LINE1 RNA expression measured by RT-qPCR upon knockdown of YTHDC1. Points represent biological replicates. P-values were calculated by two-tailed Students’s *t* test. **g**, TDP-43-RIP with RT-qPCR of LINE1. Points represent 3 individual iPSN lines. P-values were calculated by two-tailed Students’s *t* test. **h,** Co-immunoprecipitation in control iPSN lines with TDP-43 antibody or IgG control, with or without nuclease treatment. The input and precipitated proteins were immunoblotted with ZCCHC8 and TDP-43 antibodies. n=3 individual control iPSN lines. Data are mean ± s.e.m. n.s., not significant, *p < 0.05, **p < 0.01, ***p < 0.001, ****p < 0.0001, two-tailed *t* test.

Next, we examined whether the NEXT complex-mediated LINE1 RNA decay is disturbed by disease pathologies linked to C9ORF72 or TDP-43 dysfunction. In a previous study, we demonstrated that C9ORF72-ALS/FTD possess a global reduction of m^6^A level^18^. Therefore, we examined whether LINE1 RNA elevations is caused by m^6^A hypomethylation in C9ORF72-ALS/FTD iPSNs. Overexpression of the methyltransferase METTL14 or knockdown of the m^6^A demethylase FTO is sufficient to increase m^6^A modification on LINE1 RNA and reduce its level (Extended Data Fig. 3a-d). Furthermore, we examined how the NEXT complex mediates m^6^A-modified LINE1 RNA degradation. We previously found that the m^6^A reader protein YTHDC1 interacts with ZCCHC8 in neuron^18^, supporting the hypothesis that YTHDC1 directs m^6^A-marked LINE1 RNA to the NEXT complex for degradation, and reduced m^6^A on these RNAs would lead to escaped turnover. Thus, we examined the interaction between YTHDC1 and LINE1 RNA. YTHDC1-RIP coupled with RT-qPCR showed selective binding of YTHDC1 to LINE1 RNA, but not *SMARCA1* mRNA that is not m^6^A-modified^18^ (Fig. 2e). Moreover, knockdown of YTHDC1 in neurons (Extended Data Fig. 3e) increased LINE1 RNA level (Fig. 2f). These results support that the m^6^A modification on LINE1 RNA mediates its degradation through the NEXT pathway, and that the hypomethylation in C9ORF72-ALS/FTD leads to reduced decay and elevated accumulation.

There has been evidence that TDP-43 can bind LINE1 RNA directly^27,28^. We tested the possibility that TDP-43 may target its bound LINE1 RNAs to the NEXT complex for degradation. First, we confirmed the interaction between LINE1 RNA and TDP-43 in neurons using TDP-43-RIP followed by RT-qPCR (Fig. 2g). Furthermore, co-immunoprecipitation showed that TDP-43 directly interacted with ZCCHC8, and this interaction was unaffected by nuclease treatment (Fig. 2h). This reveals an alternative mechanism for RNA targeting to the NEXT complex for degradation, mediated by TDP-43 binding. Therefore, TDP-43 dysfunction failed to direct the RNA targets to the NEXT complex, resulted in reduced decay and extended half-life of LINE1 RNA (Fig. 2b), potentially accounting for its increased accumulation in neurons with TDP-43 loss of function. Altogether, these results support that dysregulation of two independent, disease-relevant regulators, m^6^A and TDP-43, converge on a shared mechanism – NEXT-mediated RNA decay – underlying the abnormal LINE1 RNA elevation, a common phenotype in C9ORF72-linked familial ALS/FTD and TDP-43 proteinopathy-associated sporadic neurodegenerative diseases.

### LINE1 RNA accumulation is associated with increased chromatin accessibility and transcription

We next explored the potential consequences of LINE1 RNA accumulation. Given its chromatin enrichment (Fig. 1a), we speculated that LINE1 RNA may affect chromatin state in neurons, similar to the effects in mESCs^12^. We first assessed global chromatin accessibility by analyzing the transposase-accessible chromatin using sequencing (ATAC-seq) data from the Answer ALS consortium. We found that the chromatin accessibility in C9ORF72-ALS/FTD iPSNs was significantly increased compared to controls, especially near transcription start sites (TSSs) (Fig. 3a). We also performed a deoxyribonuclease (DNase) I-treated terminal deoxynucleotidyl transferase-mediated deoxyuridine triphosphate nick end labeling (TUNEL) assay, a fluorescence-activated cell sorting (FACS)-based method to measure global chromatin accessibility^15,16^, and we observed consistent results of increased global chromatin openness in C9ORF72-ALS/FTD (Fig. 3b). We also performed the ATAC-seq and DNaseI-TUNEL assay in TDP-43 knockdown i^3^Neurons and observed a similar increase in global chromatin accessibility resulting from TDP-43 reduction (Fig. 3c,d).

**Fig.3:**
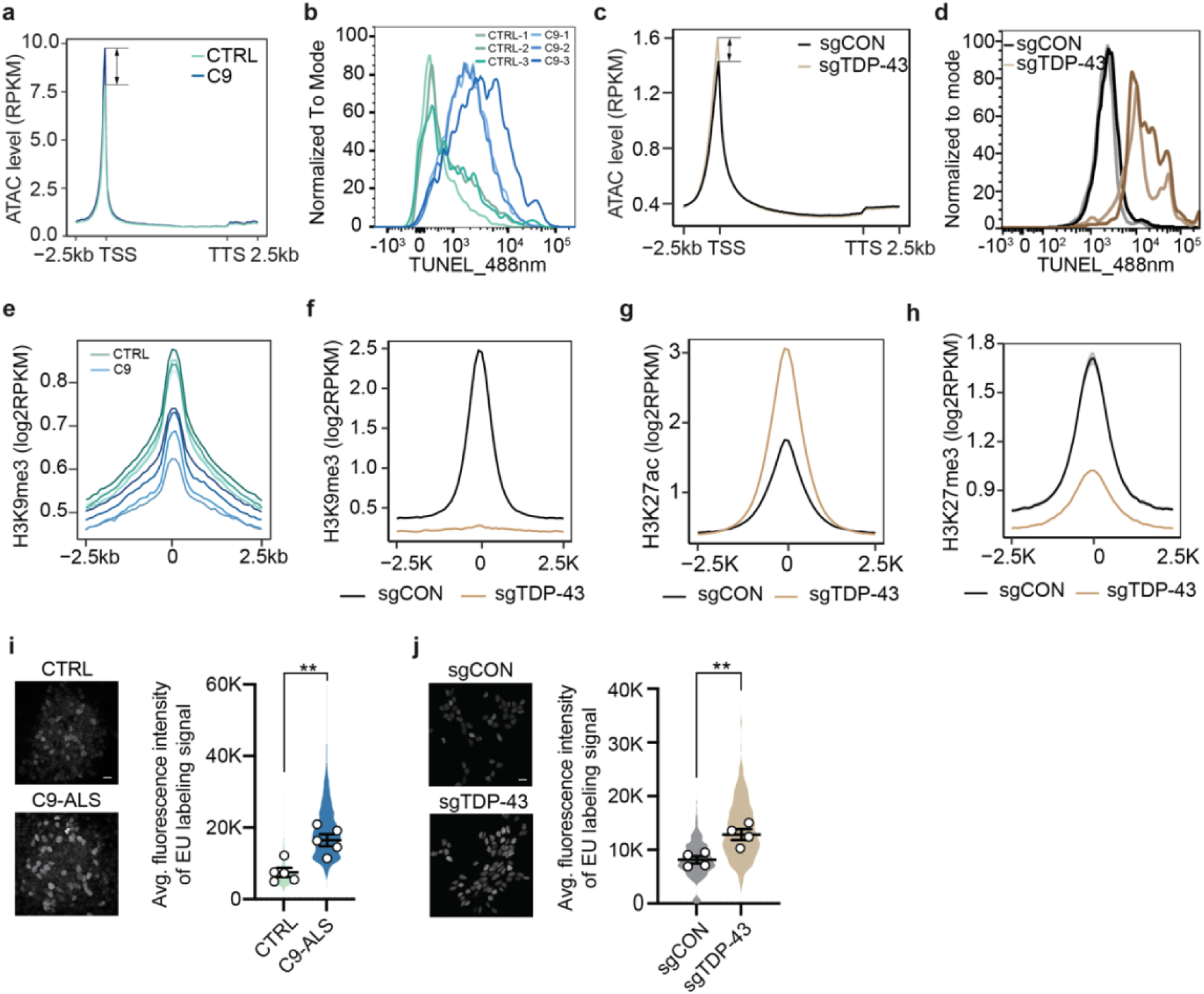
Increased chromatin accessibility and transcription in C9ORF72-ALS/FTD and TDP-43-reduced neurons. **a**, Global chromatin accessibility on gene body was quantified using ATAC-seq in control (n=42) and C9 iPSNs (n=11), from 2.5 kb upstream of the transcription start site (TSS) to 2.5 kb downstream of the transcription termination site (TTS). ATAC-seq datasets were obtained from Answer ALS consortium. **b**, Analysis of chromatin accessibility in control and patient iPSN lines by DNase I-treated TUNEL assay. The fluorescence signal was measured by flow cytometry. n=3 individual iPSN lines per group. **c,** Global chromatin accessibility on gene body was quantified using ATAC-seq in sgCON and sgTDP-43 i^3^Neurons. n=2 biological replicates per group. **d,** DNase I-treated TUNEL assay, measuring chromatin accessibility in sgCON and sgTDP-43 i^3^Neurons. n=2 biological replicates per group. **e,** Profiles of H3K9me3 level in control and patient iPSN lines (n=4 per group), centered on peak regions with ±2.5 kb flanking sequences. **f-h,** Profiles of the repressive histone marks, (f) H3K9me3 and (g) H3K27me3, and the active histone mark (h) H3K27ac, in sgCON and sgTDP-43 i^3^Neurons (n=2 per group), centered on peak regions with ±2.5 kb flanking sequences. **i-j**, Analyses of nascent RNA synthesis in (i) control and C9 iPSNs, and (j) sgCON and sgTDP-43 i^3^Neurons, measured by a click-it EU labeling imaging assay (left). Average fluorescence intensities were quantified by ImageJ (right). Scale bars, 20 µm. Points represent individual control or patient lines (n=5 per group), or sgCON or sgTDP-43 biological replicates (n=4 per group). P-values were calculated by two-tailed *t* test on the mean intensities of individual lines. Data are mean ± s.e.m. **p < 0.01, two-tailed Student’s *t* test.

We then examined the chromatin states underlying the chromatin accessibility changes potentially influenced by LINE1 accumulation. Specifically, we explored key histone modifications, including the active mark H3K27ac, and repressive marks H3K27me3 (facultative) and H3K9me3 (constitutive)^39,40^. Chromatin immunoprecipitation sequencing (ChIP-seq) revealed decreased levels of the heterochromatin mark H3K9me3 in both C9ORF72-ALS/FTD and TDP-43 knockdown neurons (Fig. 3e,f), which was further validated by immunofluorescence (IF) staining (Extended Data Fig. 4a,b). TDP-43 knockdown i^3^Neurons also exhibited increased H3K27ac and decreased H3K27me3 levels, as shown by ChIP-seq (Fig. 3g,h) and IF staining (Extended Data Fig. 4c,d), consistent with the observed increase in chromatin accessibility (Fig. 3c,d). In contrast, no changes in H3K27ac or H3K27me3 levels were detected in C9ORF72-ALS/FTD iPSNs (Extended Data Fig. 4e,f). This indicates a primary derepression of constitutive heterochromatin marks in C9ORF72-ALS/FTD, while both facultative and constitutive histone marks can be modulated by TDP-43. Finally, we validated these findings through immunostaining of postmortem brain tissues from patients. Reduced H3K9me3 levels were observed in both C9ORF72-linked ALS/FTD (Extended Data Fig. 5a) and sporadic FTD (Extended Data Fig. 5b). Moreover, increased H3K27ac and reduced H3K27me3 levels correlated with TDP-43 pathologies in sporadic FTD brain tissues (Extended Data Fig. 5c,d), highlighting specific epigenetic alterations in cells with TDP-43 dysfunction. Overall, the histone modification changes are consistent with the increased chromatin accessibility in both C9ORF72- and TDP-43-associated ALS/FTD.

Both increased chromatin openness and chromatin state changes, characterized by reduced repressive histone marks and elevated active marks, can lead to global upregulation of gene transcription. We therefore measured transcription activity using the 5-ethynyl uridine (EU)-Click iT imaging assay. In both C9ORF72-ALS/FTD iPSNs and TDP-43 knockdown i^3^Neurons, we observed a significant increase in nascent transcripts, labeled by click chemistry-marked fluorescence (Fig. 3i.j), supporting elevated transcriptional activity in both disease conditions.

We further investigated whether the epigenetic changes overlap with LINE1 loci. We observed decreased H3K9me3 at LINE1 loci in both C9ORF72-ALS/FTD (Fig. 4a) and TDP-43 knockdown neurons (Fig. 4b). In the latter, we also detected increased H3K27ac and decreased H3K27me3 at LINE1 loci (Fig. 4c), indicating a shift toward a more transcriptionally active chromatin state at these genomic regions. In C9ORF72-ALS/FTD neurons, which exhibit global m^6^A hypomethylation ^18^, the reduction in H3K9me3 correlated with decreased m^6^A on the accumulated LINE1 RNAs (Fig. 4d) and increased local LINE1 RNA expression (Fig. 4e). Additionally, genes adjacent to upregulated LINE1 RNA exhibited a trend toward increased mRNA expression (Fig. 4f), with a significant positive correlation (Fig. 4g). In TDP-43 knockdown neurons, neighboring genes of upregulated LINE1 RNA also showed significantly elevated mRNA expression (Fig. 4h), along with a strong cis correlation between LINE1 RNA levels and adjacent gene expression (Fig. 4i). Altogether, these results suggest LINE1 RNA accumulation exerts a cis-regulatory effect on local chromatin state and gene expression, likely contributing to disease-associated dysfunction and neurodegeneration.

**Fig.4:**
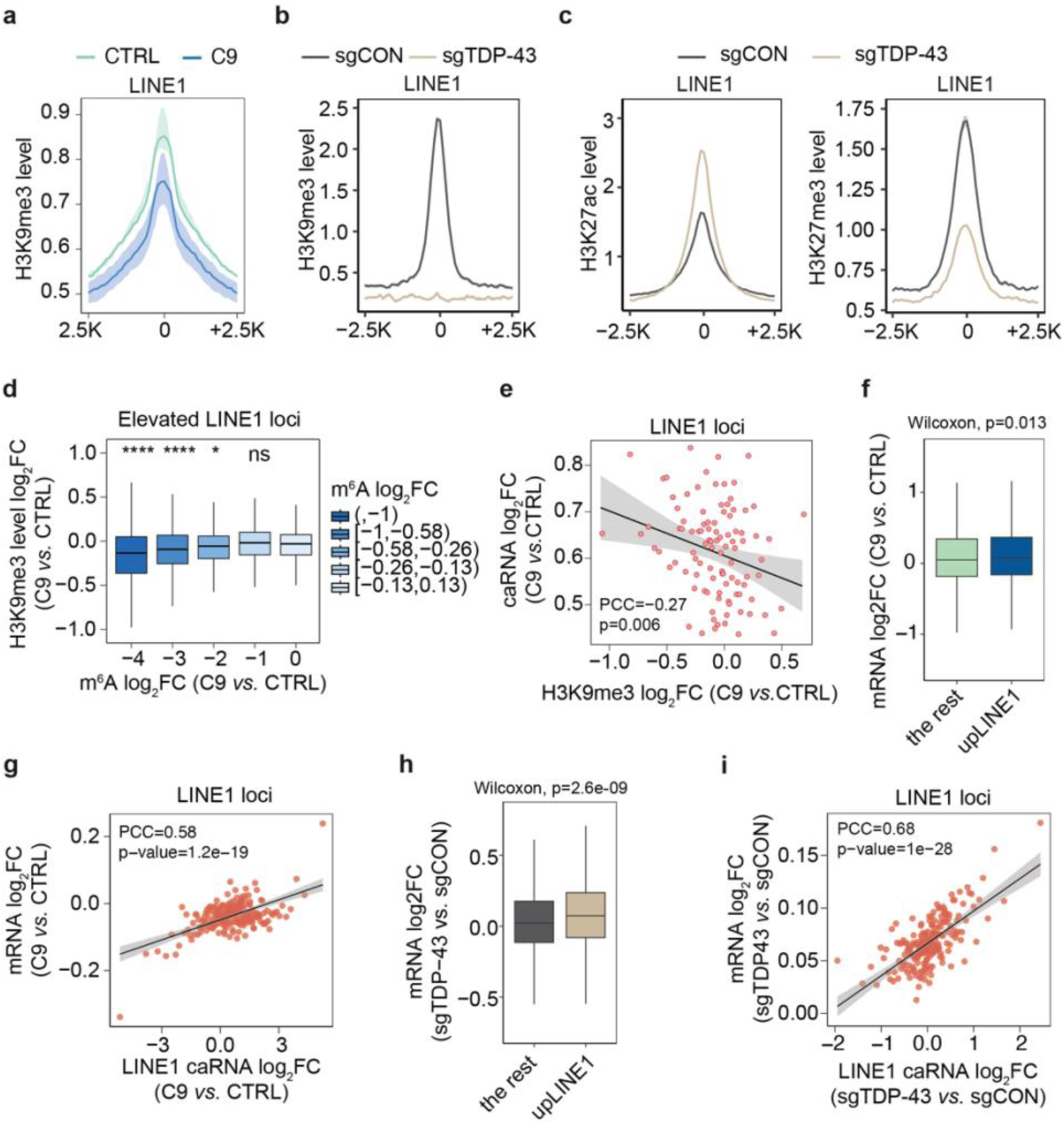
Association of LINE1 RNA expression and epigenetic changes. **A,** Profiles of H3K9me3 level at LINE1 loci. The peak center and the ±2.5 kb flanking regions were analyzed in control and patient iPSN lines (n=4 individual lines per group). **b**, Profiles of H3K9me3 at LINE1 loci with ±2.5 kb flanking regions in control and TDP-43-reduced neurons. n=2 biological replicates per group. **c,** Profiles of H3K27ac and H3K27me3 levels at LINE1 loci with ±2.5 kb flanking regions in control and TDP-43-reduced neurons. n=2 biological replicates per group. **d,** Boxplots showing positive correlation of changes between m^6^A and H3K9me3 at LINE1 loci. n=4 individual lines per group. P-values were calculated by Wilcoxon rank sum test. **e,** Scatter plot showing the negative correlation of changes in repressive histone mark H3K9me3 level and local RNA abundance at LINE1 loci, comparing patient iPSN lines with their controls. Pearson correlation coefficient (PCC) and p values are shown. **f,** Boxplots showing adjacent mRNA expression changes with or without LINE1 RNA upregulation in C9ORF72-ALS/FTD patient iPSNs (n=4 individual lines per group). **g,** Scatter plot showing the positive correlation of changes in LINE1 RNA and the expression of adjacent genes, comparing C9 patient iPSN lines with controls. Pearson correlation coefficient (PCC) and p-values are shown. **h,** Boxplots showing adjacent mRNA expression changes with or without LINE1 RNA upregulation in TDP-43-reduced neurons (n=2 biological replicates per group). **i,** Scatter plot showing the positive correlation of changes in LINE1 RNA and the expression of adjacent genes, comparing TDP-43-reduced neurons with controls. Pearson correlation coefficient (PCC) and p-values are shown.

### LINE1 ASO treatment rescues chromatin abnormality and disease related phenotypes

To delineate the causative effect of LINE1 RNA on chromatin and disease pathogenesis, we knocked down LINE1 RNA in disease-relevant neurons using a LINE1-targeting antisense oligonucleotide (ASO)^7^ (Fig. 5a), which induces RNase H1-dependent degradation of bound RNAs in the nucleus^41^. The reduction of LINE1 RNA shifted global chromatin accessibility to a more closed state in both C9ORF72-ALS/FTD iPSNs (Fig. 5b) and TDP-43-reduced neurons (Fig. 5c). Accordingly, the abnormally reduced repressive mark H3K9me3 was increased in C9ORF72-ALS/FTD iPSNs (Extended Data Fig. 6a), and the series of dysregulated histone marks H3K9me3, H3K27me3, and H3K27ac were also rescued in TDP-43 knockdown neurons (Extended Data Fig. 6b). The chromatin restoration further mitigated the globally elevated transcriptional activity (Fig. 5d,e). We also validated several targets at different genomic loci, showing that ASO-mediated LINE1 RNA reduction restored the dysregulated histone modification and decreased nascent transcripts at nearby loci (Extended Data Fig. 6c,d).

**Fig.5:**
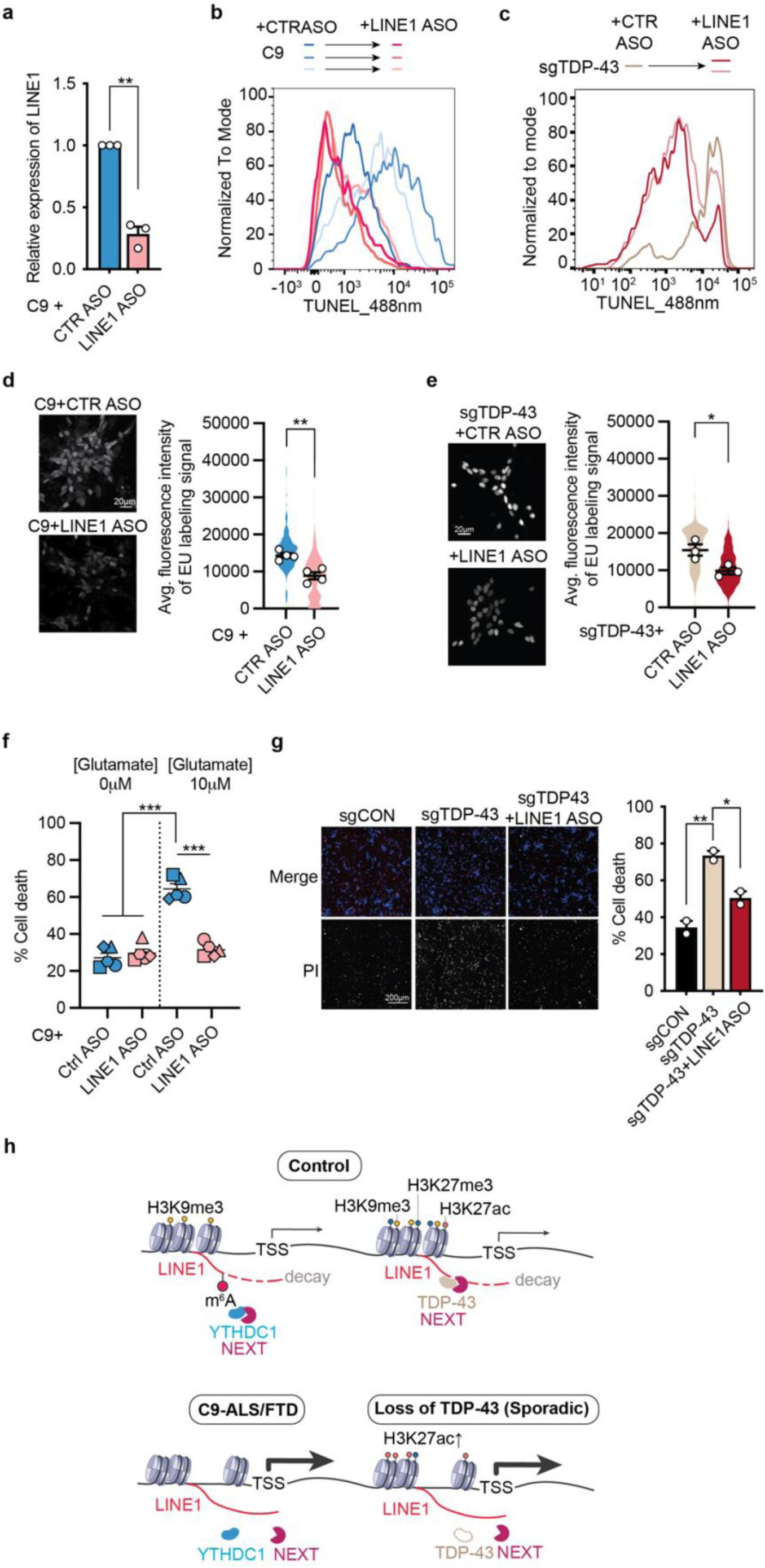
Reducing LINE1 RNA rescues chromatin and transcription abnormalities and alleviates neurodegeneration. **a,** Relative expression of LINE1 RNAs in C9 iPSN lines treated with antisense oligonucleotides (ASO) targeting LINE1 or non-targeting control. Points represent individual patient lines. P-values were calculated by paired two-tailed *t* test. **b,** Analysis of chromatin accessibility in individual patient iPSN lines following LINE1 RNA knockdown by ASO. DNase I-treated TUNEL assay was performed and analyzed by flow cytometry. n=3 individual patient lines per treatment. **c,** Analysis of chromatin accessibility in sgTDP-43 i^3^Neurons following LINE1 RNA knockdown using ASO (n=1-2), by DNase I-treated TUNEL assay. **d,** Analysis of nascent RNA synthesis in C9 iPSNs following LINE1 knockdown by ASO, measured by a click-it EU labeling imaging assay (left). Average fluorescence intensities were quantified by ImageJ (right). Scale bars, 20 µm. Points represent individual patient lines (n=4) with indicated treatment. P-values were calculated by paired two-tailed *t* test on the mean intensities of individual lines. **e,** Analysis of nascent RNA synthesis in sgTDP-43 i^3^Neurons following LINE1 knockdown by ASO, measured by a click-it EU labeling imaging assay (left), and its quantification (right). n= 3 biological replicates per treatment. Scale bars, 20 µm. **f,** Glutamate-induced excitotoxicity in C9ORF72-ALS/FTD iPSNs after treatment of LINE1 or control ASOs. Dead cells were labeled with propidium iodide (PI) and the percentage of death was quantified as the ratio of PI^+^/DAPI^+^ cells. Different shapes represent different patient iPSN lines (n=5). P-values were calculated by paired two-tailed Student’s *t* test. **g,** PI staining (left) and cell death quantification (right) of sgTDP-43 i^3^Neurons following LINE1 knockdown by ASO. Scale bars, 200 µm. **h,** A schematic model showing a convergent mechanism by which m^6^A hypomethylation in C9ORF72-ALS/FTD and loss of TDP-43, as observed in TDP-43 proteinopathy-associated sporadic neurodegenerative diseases, disrupt NEXT-mediated LINE1 RNA decay, leading to chromatin state and transcriptional dysregulation. Data are mean ± s.e.m. *p < 0.05, **p < 0.01, ***p < 0.001, paired two-tailed Student’s *t* test.

To distinguish whether the chromatin rescue resulted from LINE1 RNA or LINE1 retrotransposition, we treated neurons with azidothymidine (AZT), a compound that effectively inhibits LINE1 reverse transcriptase activity *in vitro* and *in vivo*^12,42,43^. AZT treatment did not rescue the global chromatin abnormality in patient iPSNs (Extended Data Fig. 6e), suggesting that chromatin dysregulation is mediated by LINE1 RNA and is independent of retrotransposition activity.

Finally, we examined whether the chromatin and transcriptional restorations by LINE1 RNA reduction was sufficient to ultimately improve neuronal survival phenotypes. C9ORF72-ALS/FTD patient iPSNs showed increased vulnerability to glutamate-induced excitotoxicity (Fig. 5f)^44^. Treatment of the LINE1-targeting ASO significantly improved neuron survival upon glutamate insult (Fig. 5f). TDP-43 knockdown induced progressive neuronal death. LINE1 ASO treatment also drastically increased survival of the TDP-43 deficient neurons (Fig. 5g). Altogether, these findings underscore the potential of targeting LINE1 RNA to rescue neurons under pathological conditions of C9ORF72-ALS/FTD and TDP-43 deficiency.

## Discussion

In summary, we report that LINE1 RNA is predominantly associated with chromatin and has a crucial function in maintaining the chromatin landscape and regulating chromatin-mediated gene expression in neurons. Abnormal accumulation of LINE1 RNA leads to increased chromatin accessibility and global transcriptional elevation, ultimately contributing to neurodegeneration (Fig. 5h). Our work uncovers a retrotransposition-independent RNA function of LINE1, whose dysregulation is associated with aging and multiple neurodegenerative diseases, highlighting a novel epigenetic mechanism by which LINE1 elevation contributes to disease pathogenesis.

Additionally, we elucidated the molecular mechanism underlying LINE1 RNA regulation via degradation by the NEXT pathway – a previously underappreciated post-transcriptional regulatory mechanism contributing to LINE1 elevation. Pathological changes in m^6^A modification on LINE1 RNA or in TDP-43, which can bind to this RNA, impair its targeting for NEXT-mediated degradation, resulting in abnormal LINE1 RNA accumulation (Fig. 5H). These findings highlight the critical role of RNA metabolism in neurons and provide novel insights into the molecular mechanisms of LINE1 dysregulation in neurological diseases.

This study reveals a convergent pathogenic mechanism shared between C9ORF72-associated familial ALS/FTD and TDP-43 proteinopathy-linked sporadic cases. Given that m^6^A dysregulation and TDP-43 pathology have also been found in other neurodegenerative diseases, including Alzheimer’s disease^25,45–48^, our findings suggest broader implications of this mechanism beyond ALS and FTD. It is noted that different LINE1 elements may be affected depending on the modulator, such as m^6^A methylation versus TDP-43 binding. We also observed varying histone modification changes, despite the consistent global effects on chromatin accessibility. Additional mechanisms may be involved in the epigenetic regulation, which needs further exploration. Furthermore, it is likely that LINE1 RNA-mediated chromatin remodeling enhances the transcriptional activity of additional LINE1 elements, forming a positive feedback loop that drives further LINE1 RNA accumulation and chromatin relaxation, thereby exacerbating neurotoxicity and progressive degeneration. Overall, the post-transcriptional mechanisms underlying LINE1 RNA dysregulation and its retrotransposition-independent regulatory RNA function offer a new perspective on RNA metabolism, retrotransposon elements, and neurodegeneration.

## Methods

### Plasmids

For lentiviral knockdown of *FTO* (Sigma-Aldrich, TRCN0000246247), *YTHDC1* (Sigma-Aldrich, TRCN0000243989), and *ZCCHC8* (targeting sequence: 5’-TTAGCACTGAGAGCTATTTAA-3’), and for overexpression of *METTL14* in neurons, we used previously constructed shRNA and overexpression plasmids. The design and cloning of lentiviral plasmids are as described ^18^.

The sgRNA targeting TDP-43 was cloned into pU6-control sgRNA EF1α-puro-T2A-BFP vector ^49^ (Addgene 60955). sgRNA sequence was as follows: non-targeting control: GGACTAAGCGCAAGCACCTA, and *TDP-43*: GGCCCGCGCGTGCCAGCCGA.

### Cell culture and transfection

293T cells were grown in DMEM supplemented with 10% (v/v) FBS, 100 μg/ml streptomycin and 100 U/ml penicillin. PC-3 cells were obtained from Dr. Kenneth Pienta’s lab. PC-3 cells were cultured in RPMI 1640 supplemented with 10% (v/v) FBS and 1% Penicillin-Streptomycin. All cells were maintained at 37°C with 5% CO2. 293T cells were used to package lentivirus for shRNAs, sgRNAs, and overexpression vectors as previously described ^18^.

### Human iPSC and iPSC-derived neurons

C9ORF72-ALS/FTD and non-neurological control iPSC lines were acquired from the Answer ALS repository at Cedars-Sinai iPSC Core (see Table S1 for demographics). Verification of the presence or absence of the C9ORF72 repeat expansion was conducted using repeat-primed PCR (RP-PCR)^50^. Routine assessment of poly(GP) expression level were performed on iPSCs and iPS-neurons. iPSCs were maintained in mTeSR Plus media (STEMCELL technologies, Cat# 100-0276) on growth factor reduced Matrigel (Corning, Cat# 354230), following the Cedars-Sinai Standard Operating Procedure (SOP, https://www.cedars-sinai.edu/content/dam/cedars-sinai/research/documents/biomanufacturing/complete-ipsc-culturing-protocol-rev-4-2020.pdf).

Differentiation of iPSCs into spinal neurons was achieved using the direct induced motor neuron protocol as described ^51^. The i^3^Neuron iPSCs ^52^ were maintained in Essential 8 media (Thermo Fisher Scientific, Cat# A1517001) on Matrigel (Corning, Cat# 354277) and differentiated into i^3^Neuron of Day 14 using the protocol as described ^49^. All cells were maintained at 37°C with 5% CO2.

The experiments regarding overexpression of METTL14 or knockdown of FTO in C9ORF72-ALS/FTD iPSNs were performed as previously described ^18^. In brief, neurons were transduced with lentivirus expressing GFP or METTL14, or control (Addgene, 10879) or FTO shRNA on Day 18 post-differentiation, and then harvested 7 days after infection. To harvest relatively pure neuron population from the culture, the iPSN matrix was gently washed off by 1×PBS.

For the LINE1 RNA knockdown experiment in patient iPSNs, a non-targeting control ASO (141923, 5’-CCTTCCCTGAAGGTTCCTCC-3’) ^53^ or an ASO targeting human LINE1 (5’-ACTTCCCTTCTCGCTTCATTT-3’) ^7^ was used. For chromatin accessibility assay and EU labeling assay, neurons were treated with 0.3µM ASOs on day 18 post differentiation and harvested after 7 days. For assays on specific LINE1 loci, neurons were treated with 0.5µM ASOs on day 18 post differentiation and harvested after 7 days. For glutamate-induced excitotoxicity assay, patient iPSNs were treated with 0.3µM ASOs on day 25 post differentiation, and on day 32 of differentiation, neurons were treated with 0 or 10μM L-glutamate for 4 hours to evaluate the neuron vulnerability towards glutamate-induced excitotoxicity as described ^18^. Complete media change was performed the day before ASO addition. ASOs were replenished with media change every two days.

To achieve a stable knockdown of *TDP-43* in i^3^Neurons, i^3^N-iPSCs ^52^ that constitutively express the clustered regularly interspaced short palindromic repeat inactivation (CRISPRi) module ^49^ were transduced with lentivirus expressing either an sgRNA targeting *TDP-43,* or a non-targeting control sgRNA, and the stable iPSC lines were established as previously described ^54^.

To knock down *ZCCHC8* or *YTHDC1* in i^3^Neurons, cells were transduced with lentivirus expressing the control, or *ZCCHC8* shRNA, or *YTHDC1* shRNA on differentiation day 5 at an MOI of 1.5 in Cortical Neuron Culture Medium ^49^, and harvested on differentiation day 14 for the measurement of endogenous *ZCCHC8, YTHDC1,* and LINE1 expression.

To knock down LINE1 RNA in sgTDP-43 i^3^Neurons, non-targeting control ASO and LINE1-targeting ASO were added on differentiation day 7 at 0.5 µM. Treated neurons were harvested on day 14 for all the assays. The treatment of LINE1 RT inhibitor AZT (Sigma-Aldrich, A2169) followed the same timeline as ASO treatment. AZT was used at 1 µM according to previous reports^12,42^.

### Immunoblotting

Samples were lysed in RIPA buffer supplemented with 1× cOmplete protease inhibitor cocktail (Sigma) and homogenized by shearing with 26-gauge needle for 6 times. The protein concentrations of supernatants were determined using BCA protein assay (Thermo Fisher Scientific) and measured on Tecan Infinite 200 PRO. Equal amount (20µg) of protein samples were loaded on 10% acrylamide gel and transferred to nitrocellulose membrane (Cytiva, Cat# 10600002). Membrane was blocked in 5% non-fat milk/TBST for 1 hour at room temperature and incubated with primary antibody overnight at 4 °C. After TBST wash, the membrane was incubated with HRP-linked secondary antibody (Cytiva, 1:10,000) for 1 hour. Protein bands were detected using Clarity Western ECL Substrate kit (Bio-Rad) and imaged using a ChemiDoc Imager (Bio-Rad). Primary antibodies: anti-LINE1 ORF1p antibody, clone 4H1 (1:500, Millipore Sigma MABC1152), anti-GAPDH antibody (1:1000, Cell Signaling Technology #2118), anti-TDP-43 (1:1000, Proteintech 10782-2-AP), anti-α-tubulin (1:2000, Thermo Fisher 62204), anti-histone H3 (1:1000, Cell Signaling Technology #4499), anti-Coilin (1:1000, Abcam, ab87913), anti-TUJ1 (1:1000, Cell Signaling Technology #4466S), and ZCCHC8 (1:1000, Proteintech 23374-1-AP). Experiments were repeated three times.

### Immunoprecipitation

One 10-cm dish of iPSNs was harvested on differentiation day 18. Neurons were washed twice with PBS, the the neuron matrix was washed off by cold PBS. The immunoprecipitation procedure followed the protocol as described ^18^. After cell lysis, protein concentrations were measured by BCA assay. 1mg lysate was used per pull-down. For RNase-treated lysates, 1ul of RNase cocktail enzyme mix (Thermo Fisher AM2286), 1ul of Benzonase nuclease (Millipore E1014-25KU), and MgCl_2_ (3mM final concentration) were added to the lysate and incubated at 30°C for 30min. Immunoprecipitation was performed using a ratio of 5µg antibody, 30ul protein G Dynabeads (Invitrogen 10004D), and 1mg lysate. Antibodies for immunoprecipitation: anti-TDP-43 (Proteintech 10782-2-AP) and anti-rabbit IgG (Sigma-Aldrich). Antibodies for immunoblotting: anti-TDP-43 (1:1000, Abnova H00023435-M01) and anti-ZCCHC8 (1:1000, Proteintech 23374-1-AP). The experiment was repeated with 4 individual control iPSN lines independently.

### Immunofluorescence

Neurons were cultured on Matrigel-coated coverslips in 24-well plates. Cells were fixed with 4% PFA/PBS for 15min at room temperature, permeabilized with 0.2% Triton X-100/PBS for 5min, and blocked in blocking buffer (1% BSA, 2% goat serum, 1× PBS) for 30min. Samples were incubated with a primary antibody overnight at 4 °C, and further incubated with Alexa Fluor 488/546-conjugated secondary antibodies (Thermo Fisher Scientific, 1:500). The nuclei were stained with DAPI. Cells were imaged using Zeiss LSM 900. The intensity of signal was quantified by Image J. Primary antibodies: H3K9me3 (1:1000, Abcam ab8898), H3K27me3 (1:1000, Sigma-Aldrich 07-449), H3K27ac (1:1000, Active Motif 39133) and TDP-43 (1:1000, Abnova H00023435-M01).

### DNase I treated-TUNEL assay

Two million neurons were harvested by PBS. The cells were permeabilized in 0.5% Triton X-100 in PBS buffer for 15 min, and then treated with 0.5 U/ml of RNase-free DNase I (Promega, Cat# M6101) for 5 min. After DNase treatment, a fluorescent TUNEL assay was performed using the DeadEnd Fluorometric TUNEL System (Promega, Cat# G3250), following the manufacturer’s protocol. The fluorescent signals were measured by flow cytometry (BD LSRII), and data was analyzed using Flowjo v10.4.

### RNA isolation and qRT-PCR

Total RNAs were isolated with TRIZOL. RNA samples used for RT-qPCR were treated with RQ1 DNase I (Promega) and then used for complementary DNA (cDNA) synthesis with random hexamers (Thermo Fisher Scientific). qPCR was performed using PowerUp SYBR Green Master Mix (Thermo Fisher Scientific, Cat# A25776) and Taq Pro Universal SYBR qPCR master mix (Vazyme, Cat# Q712-03) on the QuantStudio 6 real-time PCR system (Thermo Fisher Scientific). At least three biological replicates with each containing two technical replicates were used for quantification. *HPRT1* mRNA was used as the internal control. Inter-group differences were assessed by two-tailed Student’s *t*-test.

### MeRIP-seq and MeRIP-RT-qPCR

Ribosomal RNA was depleted from chromosome-associated RNA by RiboMinus Eukaryote Kit v2 (Thermo Fisher Scientific). m^6^A-RNA immunoprecipitation (MeRIP) was performed using the EpiMark *N*^6^-Methyladenosine Enrichment Kit (New England Biolabs, Cat# E1610S) following the manufacturer’s protocol. RNAs from input and MeRIP were purified with RNA clean and concentrator-5 (Zymo, Cat# R1014) for either library preparation for MeRIP-seq or MeRIP-RT-qPCR. Library was prepared using the SMARTer Stranded Total RNA-Seq Kit v2 (Takara) and sequenced on a NovaSeq system with paired-end 150bp reads. For MeRIP-RT-qPCR, the RNAs were subjected to reverse transcription and qPCR. The *GLuc* spike-in RNA was used as normalization control. The fold enrichment of MeRIP was compared to the input. Inter-group differences were assessed by two-tailed Student’s *t*-test.

### RNA decay measurement

5 μg/ml Actinomycin D was added to day-32 iPSNs and day-14 i^3^Neurons at 6 h, 3 h, and 0 h before collection for RNA extraction by Trizol. To measure the half-life of LINE1 RNA, equal amount of *CLuc* RNA (New England Biolabs, Cat# E1610S) was spiked in before reverse transcription as normalization control for qPCR.

### Nascent RNA measurement

For fluorescence labeling assay, neurons were cultured on Matrigel-coated coverslips in 24-well plates. The basal nascent RNA measurements were performed on control or patient iPSNs of differentiation day 32, or sgCON and sgTDP-43 i^3^Neurons of differentiation day 14. The day before the assay, a complete media change was performed. The nascent RNA labeling assay followed the manufacturer’s protocol of Click-iT RNA Alexa Fluor 488 Imaging Kit (Invitrogen C10329). An equal volume of 2× working solution of 5-Ethynyl Uridine (EU) was prepared in pre-warmed complete media and added to the cell media to achieve a 1× final working solution. Cells were incubated with EU for 1 hour in the cell culture incubator. After incubation, cells were immediately fixed by 3.7% formaldehyde and proceeded to Click-iT reaction. Hoechst 33342 was used to stain the nuclei and the coverslips were then mounted for imaging. Cells were imaged using Zeiss LSM 900, and the intensity of signal was quantified by Image J.

For nascent transcript quantification by RT-qPCR, one million cells were seeded to enter the terminal neuron maturation in 6-well plates. The same treatment timeline was used as the nascent RNA imaging assay. Cells were incubated with EU for 1 hour. Total RNA was purified by RNA Clean & Concentrator-5 (Zymo) with on-column DNase treatment to remove potential DNA contaminants. Nascent RNA capture was performed following the manufacture’s manual of Click-iT Nascent RNA Capture Kit (Invitrogen, Cat# C10365) with some modifications. 2ng of EU-labeled spike-in *luciferase* RNA was added to 1µg of EU-labeled RNA for biotinylation and pull-down. cDNA synthesis was performed on beads and eluted by heating at 85 °C in SuperScript IV VILO master mix (Invitrogen, Cat# 11756050). The *luciferase* RNA was used as normalization control. Inter-group treatment differences were assessed by paired Student’s *t*-test.

### RNA immunoprecipitation (RIP)

One 10cm dish per line of control iPSNs were harvested on day 20 of differentiation. The cell pallets were lysed in equal volume of RIP lysis buffer (50mM HEPES, 150mM KCl, 2mM EDTA, 0.5% NP40, 0.5mM DTT, supplemented with 2× cOmplete protease inhibitor cocktail (Sigma, Cat# 11697498001), 0.25μl/100μl SUPERase-in RNase Inhibitor (Thermo Fisher Scientific)), sheared with 26-gauge needle for 6 times, incubated on ice for 5min, and centrifuged at 14,000g for 30min at 4 °C. The 1:1 blend Protein A/G beads (Thermo Fisher Scientific, Cat# 10002D and 10004D) were washed once with RIP lysis buffer, and incubated with YTHDC1 (Abcam ab264375), TDP-43 (Proteintech 10782-2-AP), or ZCCHC8 (Proteintech 23374-1-AP) antibody, or rabbit IgG (Sigma) for 1 hour at room temperature. The antibody-coated beads were washed three times with RIP lysis buffer. The supernatants of the lysates were added to the coated beads at equal amounts and incubated at 4 °C with rotation for 4 hours. 5% of the lysates was saved as input and extracted by TRIZOL for RNA. The beads were washed 6 times with NT2 buffer (50mM HEPES, 200mM NaCl, 2mM EDTA, 0.05% NP40, 0.5mM DTT, 1× cOmplete protease inhibitor and RNase inhibitor). After washing, TRIZOL was added directly to the beads to elute the RNA. RNAs recovered from RIP and input were subjected to reverse transcription and qPCR. The fold enrichment of RIP was compared to the input. Inter-group differences were assessed by two-tailed Student’s *t*-test. Rabbit IgG-RIP was used as the negative control.

### Immunohistochemistry (IHC) and co-immunofluorescence (co-IF) on human tissue

Paraffin-embedded brain sections from C9ORF72-ALS/FTD patients, sporadic FTD patients with TDP-43 proteinopathies, and non-neurological controls were obtained from Johns Hopkins Brain Resource Center, Johns Hopkins and Temple University ALS Postmortem Tissue Cores (see Table S2 and Table S3 for demographic information). Tissue sections were gradually rehydrated with xylene, 100% ethanol, 95% ethanol, 70% ethanol, 50% ethanol, 30% ethanol, 1× PBS, and dH2O. For IHC, Antigen retrieval was performed with 0.1M sodium citrate buffer (pH 6.0) for 5min in a steam sterilizer. The slides were permeabilized with 0.2% Triton X-100 for 20min and blocked with the blocking buffer (1% BSA and 2% Goat Serum diluted in 1× PBS) for 1 hour. Slides were incubated with anti-H3K9me3 antibody (1:2000, Abcam ab8898) diluted in the blocking buffer overnight at 4°C. Slides were washed three times with 1× PBS, treated with 0.3% H_2_O_2_ for 30min, and incubated with Goat Anti-Rabbit IgG (H+L) biotinylated secondary antibody (Vector Laboratories BA-1000-1.5, 1:300) for 1 hour. Next, the slides were treated with the R.T.U. VECTASTAIN Elite ABC Reagent for 30min and developed using the DAB Peroxidase Substrate Kit (Vector Laboratories SK-4100). Slides were imaged with Keyence BZ-X all-in-one fluorescent microscope. All images were acquired at the same brain layer, using identical exposure time and conditions.

For co-IF of TDP-43 and histone marks on patient slides, after de-paraffinizing and rehydrating, antigen retrieval was performed using Tris-based antigen unmasking solution (Vector laboratories, H-3301-250) at 121°C for 20min. The slides were washed and permeabilized with 0.2% Triton X-100 for 15min and blocked with the blocking buffer (20% goat serum and 1% BSA diluted in 1× PBS) for 1 hour. Slides were co-incubated with mouse anti-TDP-43 antibody (1:800, Abnova H00023435-M01) and rabbit anti-H3K27me3 antibody (1:1000, Sigma-Aldrich 07-449), or rabbit anti-H3K27ac antibody (1:1000, Active Motif 39133), in the blocking buffer (5% goat serum, 1% BSA, and 0.002% Triton X-100) overnight at 4°C. The next day, slides were incubated with anti-mouse and anti-rabbit Alexa Fluor 488/546-conjugated secondary antibodies (Thermo Fisher A-11001 and A-11035), for 2 hours at room temperature (RT), and then incubated with 1% Sudan Black in 70% ethanol for 30min at RT. Sudan Black was washed away by 70% ethanol rinse. The cell nuclei were stained by DAPI. Mounted slides were imaged with Zeiss LSM 900.

### Chromatin-associated RNA isolation and RNA-seq

iPSNs were fractionated following established protocols ^16,55^. In brief, 6×10^6^-1×10^7^ neurons were harvested by 1× PBS and washed with cold PBS/1mM EDTA. To lyse the cytoplasmic/plasma membrane, cold NP-40 lysis buffer (10mM Tris-HCl pH 7.5, 0.05% NP-40, 150mM NaCl) was used to resuspend the washed cell pallet and incubated on ice for 5min. The cell lysate was then over layered over 2.5× volume of pre-chilled sucrose cushion buffer (24% RNase-free sucrose in NP-40 lysis buffer), and centrifuged for 10min at 4 °C at 15,000g. To lyse the nuclear membrane, the crude nuclei pellet was washed with cold PBS containing 1mM EDTA, resuspended in pre-chilled glycerol buffer (20mM Tris-HCl pH 8.0, 75mM NaCl, 0.5mM EDTA, 0.85mM DTT, 50% glycerol), supplemented with 1:1 volume of pre-chilled nuclei lysis buffer (10mM HEPES pH 8.0, 1mM DTT, 7.5mM MgCl_2_, 0.2mM EDTA, 0.3M NaCl, 1M urea, 1% NP-40), and incubated on ice for 2 min before centrifugation at 15,000g for 2 min at 4 °C. Finally, the pallet was gently rinsed by cold PBS/1mM EDTA and treated with RNase-free DNase (Promega, 20U) for 10 min at 37 °C to solubilize the chromatin. To evaluate the cell fraction efficiency, 10% of each fraction lysate was saved for western blotting. For chromatin-associated RNA-seq, 1ml TRIzol (Invitrogen, Cat# 15596018) was directly added to the chromatin fraction to extract chromatin-associated RNA. Isolated chromatin-associated RNAs were subject to MeRIP-seq (see below) or direct RNA-seq. RNA-seq libraries were prepared using the SMARTer Stranded Total RNA-Seq Kit v2 (Takara) and sequenced on a NovaSeq system with paired-end 150bp reads.

### Chromatin-associated RNA MeRIP-seq analysis

Raw reads were initially processed for trimming using Trimmomatic-0.39 ^56^, followed by alignment to the human genome and transcriptome (hg38, version 29, 2018-08-30), together with spike-in genomes including unmodified control RNA (Cypridina Luciferase) and m^6^A methylated control RNA (Gaussia Luciferase) (New England Biolabs) using HISAT (version 2.1.0) with ‘--rna-strandness RF’ parameters. Annotation files (version v29, 2018-08-30, in gtf format) were obtained from the GENCODE database (https://www.gencodegenes.org/). Mapped reads were separated by strands using samtools (version 1.9), and m^6^A peaks on each strand were identified with MACS (version 2)^57^, using the settings ‘-g 2.052e9--nomodel’ and ‘--keep-dup 5’ separately. Peaks with a q-value below 0.01 were considered significant. Significant peaks that were consistently identified in at least two biological replicates were merged using bedtools (v.2.26.0)^58^ and utilized for further analysis. The number of reads mapped to human genome divided by number of reads mapped to m^6^A modified spike-in represented whole m^6^A level. Functional enrichment analysis was performed with DAVID ^59^ with default parameter.

Repeat annotations, sourced from RepeatMasker (2019-01-29), were obtained from the UCSC Table Browser. Enhancer RNA (eRNA), promoter-associated RNA (paRNA), and repeat RNA were classified as RNAs derived from enhancers, promoters, and repetitive elements, respectively. To minimize noise from nascent pre-mRNA in our analysis, paRNAs were further divided into sense (paRNA [+]; coding strand) and antisense (paRNA [-]; template strand) categories based on the origin of the transcripts. The featureCounts function in R was employed to quantify reads associated with carRNAs, including eRNA, paRNA, and repeat RNA.

### Chromatin-immunoprecipitation (ChIP)-seq and ChIP-qPCR

Histone ChIP was performed following the ENCODE protocol (https://www.encodeproject.org/documents/be2a0f12-af38-430c-8f2d-57953baab5f5/@@download/attachment/Epigenomics_Alternative_Mag_Bead_ChIP_Protocol_ *v1.1_exp.pdf*) with some modifications on sonication. In brief, iPSNs at differentiation day 32 were washed three times with 1× PBS, and the neuron matrix was collected and resuspended in 1× PBS.

Neurons were cross-linked in 1% formaldehyde for 10min at room temperature and sonicated on Covaris E220 for 8-12min depending on the DNA size post sonication, following the manufacture’s protocol of truChIP Chromatin Shearing Kit (Covaris) for suspension cells. 1% of the DNA post sonication were de-crosslinked and extracted by MinElute Reaction Cleanup Kit (Qiagen, Cat# 28204) to predict the DNA amount in the rest of samples. Equal amounts of DNA from each iPSN lines were diluted at 1:1 ratio with 2× Covaris dilution buffer and incubated with anti-H3K9me3 (Abcam ab8898) or rabbit IgG antibody (Sigma) at 4 °C overnight. 5% input was saved from each sample. The immunocomplexes were recovered by incubating with Protein A/G (50/50) magnetic beads for 1 hour at 4 °C. The beads were sequentially washed with RIPA/140mM NaCl buffer (0.1% DOC, 0.1% SDS, 1% Triton X-100, 140mM NaCl, 1mM EDTA, 20mM Tris-HCl pH 8.1) twice, RIPA/500mM NaCl buffer (0.1% DOC, 0.1% SDS, 1% Triton X-100, 500mM NaCl, 1mM EDTA, 20mM Tris-HCl pH 8.1) twice, LiCl buffer twice (0.25M LiCl,1% NP40, 1% Na-Deoxycholate, 1mM EDTA, 10mM Tris-HCl pH 8.1), and TE buffer twice. All buffers were supplemented with 2× protease inhibitor. Input and ChIP fractions were eluted in ChIP elution buffer (10mM Tris-Cl pH 8.0, 5mM EDTA, 300mM NaCl, 0.1% SDS) and reverse crosslinked (reverse cross-linking buffer: 250mM Tris-HCl pH 6.5, 1.25M NaCl, 62.5mM EDTA, 5mg/ml Proteinase K, 62.5µg/ml RNAse A) at 65 °C overnight. DNAs were purified by phenol-chloroform extraction for either qPCR or high-throughput sequencing library preparation. Libraries were prepared using NEBNext Ultra II DNA library prep kit for Illumina (New England Biolabs) and sequenced on a NovaSeq system with paired-end 150bp reads.

### Chromatin accessibility profiling by ATAC-seq

ATAC-seq was performed following Kaestner Lab’s omni-ATAC-seq protocol (https://www.med.upenn.edu/kaestnerlab/assets/user-content/documents/ATAC-seq-Protocol-(Omni)-Kaestner-Lab.pdf). In brief, 50,000 neurons were harvested by cold PBS wash-off, lysed by pre-chilled resuspension buffer (10mM Tris-HCl, pH 7.5, 10mM NaCl, 3mM MgCl_2_) supplemented with 0.1% NP-40, 0.1% Tween-20, and 0.01% Digitonin, on ice for 3 minutes. Nuclei were pelleted by centrifuge. For transposition, ∼50,000 nuclei were incubated with Illumina Tagment DNA buffer and Tn5 transposase, supplemented with 0.1% Tween-20 and 0.01% Digitonin, for 30 minutes at 37 °C on a shaking thermomixer. After incubation, DNAs were purified by Qiagen MinElute Cleanup kit and proceeded with PCR amplification for library generation. The ATAC-seq libraries were sequenced on a NovaSeq system with paired-end 150bp reads.

### DNA-seq analysis

Raw reads were trimmed with Trimmomatic-0.39 ^56^ and then mapped to human genome (hg38) using bowtie2 (version 2.4.1) ^60^ with “-dovetail” parameter in default mode: search for the multiple alignments, report the best one. For ATAC-seq, reads mapped to the mitochondrial genome were excluded, and duplicated reads were removed using the Picard tool (MarkDuplicates) (http://broadinstitute.github.io/picard/). Similarly, for ChIP-seq, duplicated reads were also removed with the Picard tool (MarkDuplicates).

#### Regulatory region definition

Promoters were identified as regions extending 1 kb upstream and 100 bp downstream of the transcription start site (TSS), while enhancers were identified as H3K27ac peaks located distally from promoters, ranging from 5 kb to 50 kb away from the TSS. H3K27ac peaks were identified in iPSN cells using H3K27ac ChIP-seq data processed with HOMER^61^.

### Statistics and reproducibility

Data shown were from at least three independent biological replicates or individual control or patient cell lines. Quantitative data were presented as mean with standard error of the mean (SEM). R or Prism (GraphPad Software) were used for statistical analysis, including Student’s *t* test, one-way ANOVA with multiple testing, Wilcoxon rank sum test where appropriate. Statistical significance defined as p<0.05 (∗), p<0.01 (∗∗), p<0.001 (∗∗∗), and p < 0.0001 (∗∗∗∗). Statistical details of experiments can be found in the figure legends.

## Data and code availability

Requests for further information or resources and reagents should be directed to and will be fullfilled by the lead contact, Shuying Sun. Plasmids and cell lines generated in this study are available from the lead contact upon request.

ATAC-seq datasets from Answer ALS (https://www.answerals.org) were downloaded and analyzed in this study. The m^6^A-seq and ChIP-seq data generated by this study have been deposited in the GEO database under the accession number GSEXXXXX are publicly available as of the date of publication.

## Acknowledgments

We thank John Hopkins Brain Resource Center, Johns Hopkins, and Temple University ALS Postmortem Tissue Cores for providing postmortem human brain tissue samples. We thank Dr. Lyle W. Ostrow from Lewis Katz School of Medicine at Temple University for guidance and for providing the human tissue samples. We thank Dr. Terri Thompson from Answer ALS consortium for providing help and detailed data information. We thank Dr. Kevin Talbot at University of Oxford for providing us the isogenic pair of patient iPSC lines. We thank Dr. Michael Ward for sharing the i^3^N iPSC line and i^3^Neuron differentiation protocols. We thank Dr. Kenneth Pienta’s lab for sharing PC-3 cells. We thank Alexa Wade and Dr. Anastasia Kralli for sharing RNA spike-ins. We thank Dr. Kathleen Burns for sharing the original house-made ORF1p antibody and providing valuable suggestions. We thank Sun lab members for helpful discussion.

## Funding

National Institutes of Health grant RF1NS113820 (SS)

National Institutes of Health grant R01AG078948 (SS)

National Institutes of Health grant RF1NS127925 (SS)

National Institutes of Health grant RM1HG008935 (CH)

National Institutes of Health grant R01ES030546 (CH)

The Packard Center for ALS Research (SS)

Target ALS Springboard Postdoc Fellowship (YL)

ALS Association Milton Safenowitz Post-Doctoral Fellowship (ZZ)

Muscular Dystrophy Association Development Grant (ZZ)

Toffler Scholar Award (ZZ, YY)

The Howard Hughes Medical Institute (CH).

## Author contributions

Conceptualization: YL, XD, CH, SS

Methodology: YL, XD, CH, SS

Investigation: YL, XD

Data curation: YL, XD

Visualization: YL, XD

Funding acquisition: CH, SS

Project administration: YL, XD, YX, ZZ, YY, NW, CL

Supervision: CH, SS

Resources: KC, JCT

Writing – original draft: YL, XD

Writing – review & editing: CH, SS

## Ethics Declarations

C.H. is a scientific founder, a member of the scientific advisory board and equity holder of Aferna Bio, Inc. and Ellis Bio Inc., a scientific cofounder and equity holder of Accent Therapeutics, Inc., and a member of the scientific advisory board of Rona Therapeutics and Element Biosciences.

## Data and materials availability

Requests for further information or resources and reagents should be directed to and will be fullfilled by the lead contact, Shuying Sun. Plasmids generated in this study are available from the lead contact upon request.

