## Supplementary material for "LINE1 RNA dysregulation impairs chromatin accessibility in C9ORF72- and TDP-43-linked ALS/FTD"

### Extended Data

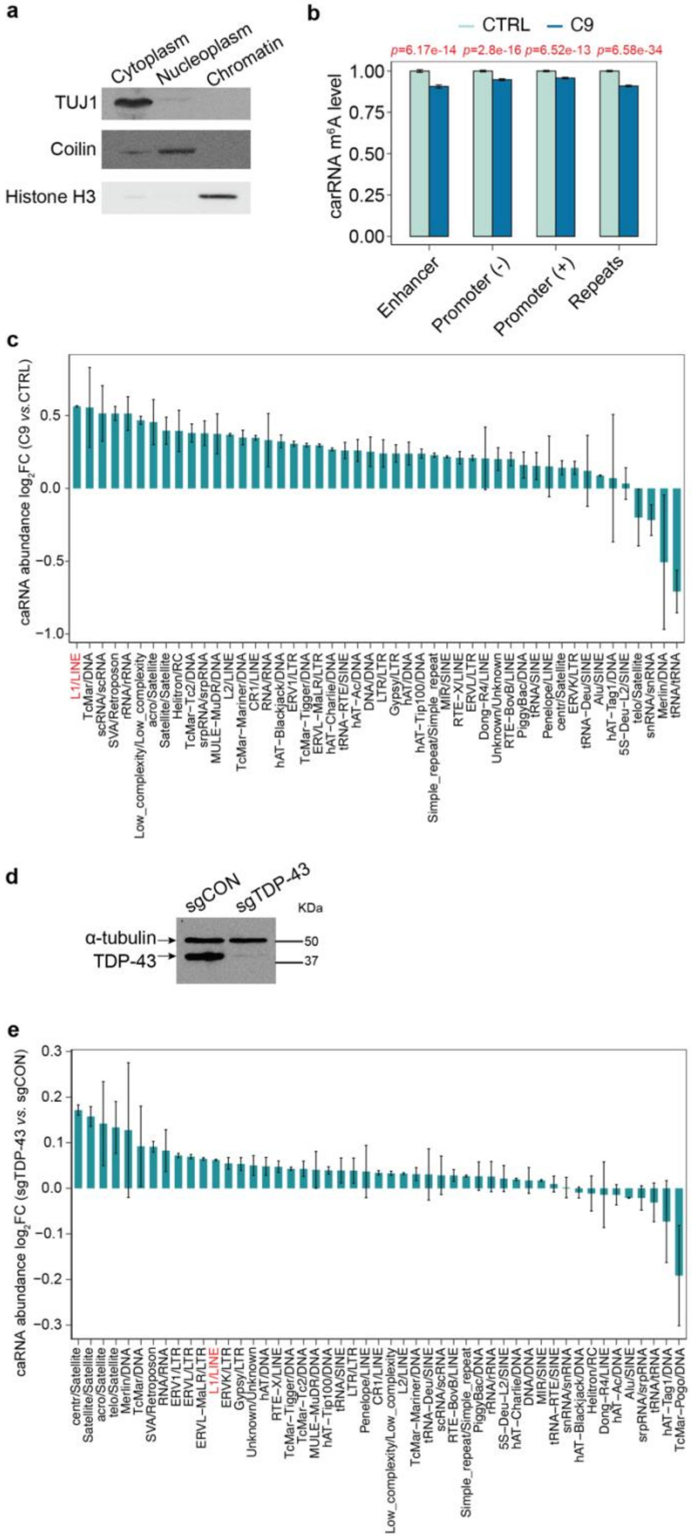

**Extended Data Fig. 1: Chromatin-associated repeat RNAs are dysregulated in C9ORF72-ALS/FTD iPSNs and TDP-43-reduced neurons.**

**a**, Cell fractionation validation shown by Western blotting analysis. **b,** Relative $m^6A$ level changes on carRNAs in control (n=4) and C9 (n=4) iPSNs, quantified through normalizing $m^6A$ sequencing results with spike-in. P-values were calculated by two-tailed Student's *t* test. **c and e,** Repeat families (x-axis) ranked by expression fold changes (y-axis), comparing (c) control and patient iPSN lines (n=4 per group) and (e) sgControl and sgTDP-43 $i^3$Neurons (n=3 per group). **d,** Western blot showing TDP-43 knockdown in TDP-43-reduced neurons (sgTDP-43). α-tubulin was used as an internal loading control.

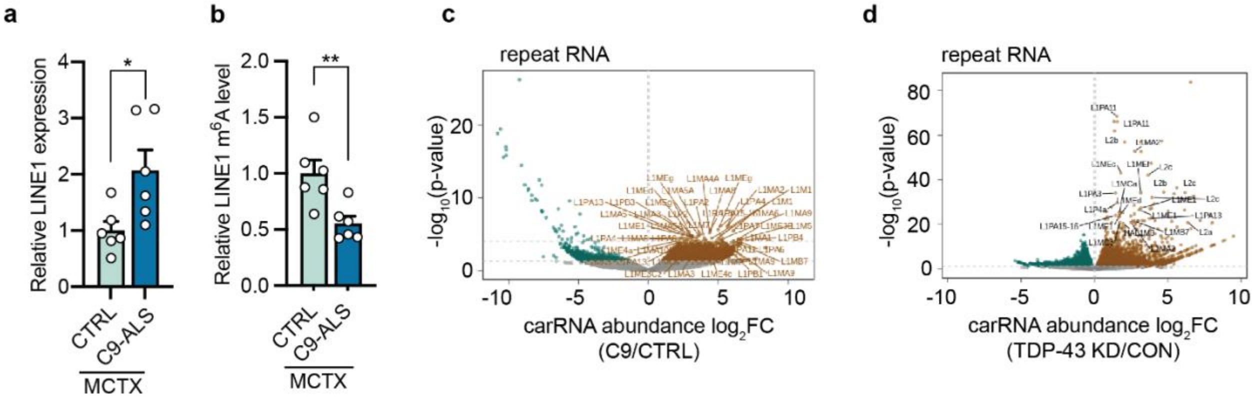


**Extended Data Fig. 2: LINE1 RNA dysregulation.**

**a**, Relative LINE1 RNA expression level measured by RT-qPCR in postmortem motor cortex. Points represent individual control or patient tissue samples (*n*=6 per group). P-values were calculated by two-tailed *t* test. **b,** Relative m$^6$A levels measured by MeRIP-RT-qPCR of LINE1 RNA in postmortem motor cortex. Points represent individual control or patient tissue samples (*n*=6 per group). P-values were calculated by two-tailed *t* test. **c-d,** Volcano plot of differentially expressed repeat RNAs in (c) C9ORF72-ALS/FTD and (d) upon TDP-43 knockdown. Upregulated LINE1 elements with $p < 0.0001$ are labeled. Data are mean ± s.e.m. *$p < 0.05$, **$p < 0.01$, two-tailed *t* test.

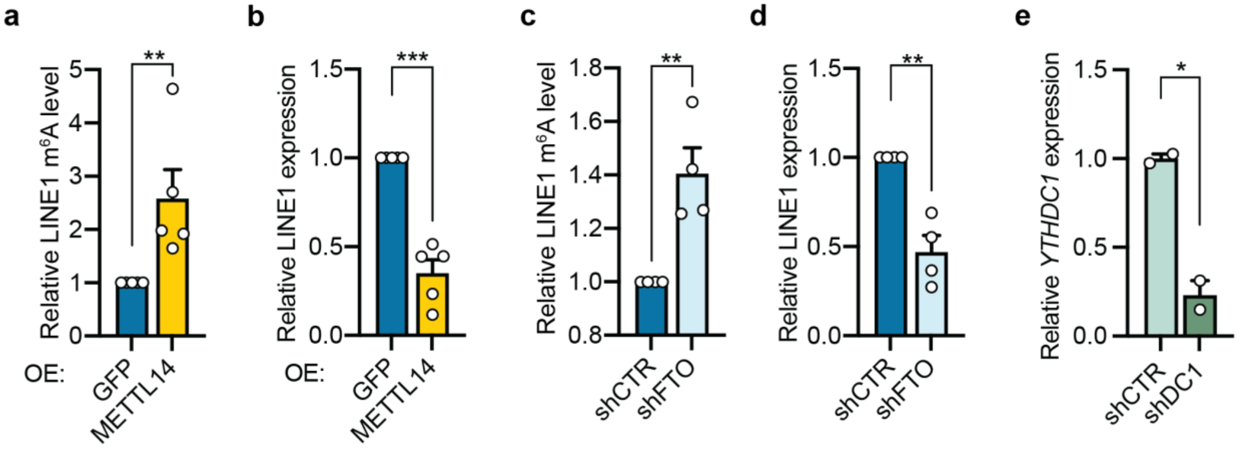


**Extended Data Fig. 3: m$^6$A regulates LINE1 RNA.**

**a-d,** The m$^6$A level measured by MeRIP-RT-qPCR (a and c) and the relative RNA expression quantified by RT-qPCR (b and d) of LINE1 in iPSNs with (a and b) overexpression of METTL14 or (c and d) knockdown of FTO. P-values were calculated by two-tailed Student's *t* test. **e**, RT-qPCR showing YTHDC1 knockdown in neurons (n=2 biological replicates per group). P-values were calculated by two-tailed *t* test. Data are mean ± s.e.m. *p < 0.05, **p < 0.01, ***p < 0.001, two-tailed *t* test.

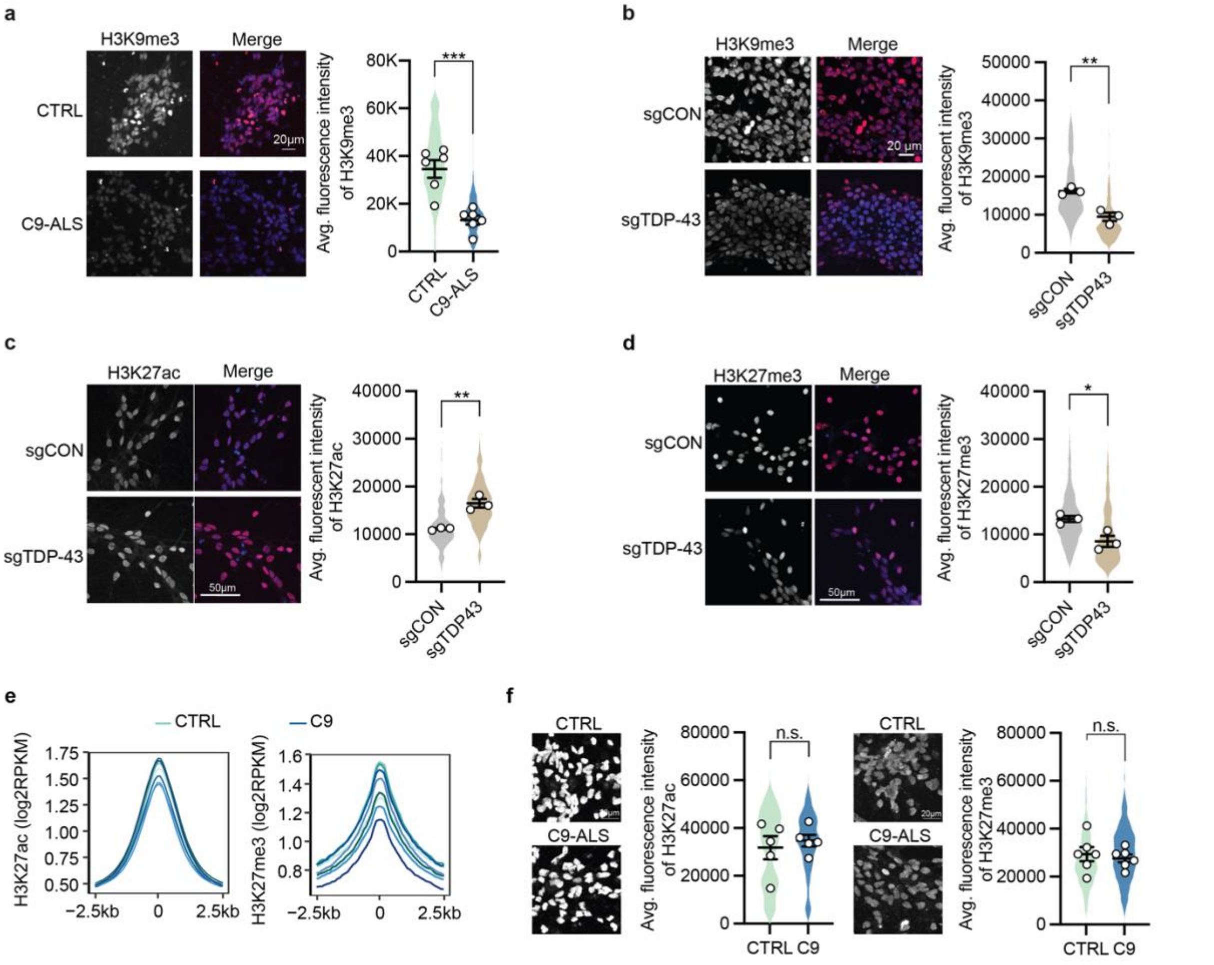


**Extended Data Fig. 4: Chromatin state changes in C9ORF72-ALS/FTD iPSNs and TDP-43-reduced neurons.**

**a,** Immunofluorescent (IF) staining and quantification of H3K9me3 expression in control and patient iPSNs. Blue, DAPI; red, H3K9me3. Points represent individual control or patient lines (n=6 per group). P-values were calculated by two-tailed *t* test on the mean intensities of individual lines. **b,** IF staining and quantification of H3K9me3 expression in control and TDP-43-reduced neurons. Blue, DAPI, red, H3K9me3. Points represent biological replicates (n=3 per group). P-values were calculated by two-tailed *t* test on the mean intensities of replicates. **c-d,** IF staining and quantification of (c) H3K27ac and (d) H3K27me3 expression in control and TDP-43-reduced neurons. Blue, DAPI, red, H3K27ac or H3K27me3. Points represent biological replicates (n=3 per group). P-values were calculated by two-tailed *t* test on the mean intensities of replicates. **e,** Profiles of H3K27ac (left) and H3K27me3 (right) levels in control and patient iPSN lines (n=4 per group), centered on peak regions with ±2.5 kb flanking sequences. **f,** IF staining and quantification of H3K27ac (left panel) and H3K27me3 (right panel) expression in control and patient iPSNs. Points represent individual control or patient lines (n=5-6 per group). P-values were calculated by two-tailed *t* test on the mean intensities of individual lines. Data are mean ± s.e.m. n.s., not significant, $*p < 0.05$, $**p < 0.01$, $***p < 0.001$, two-tailed *t* test.

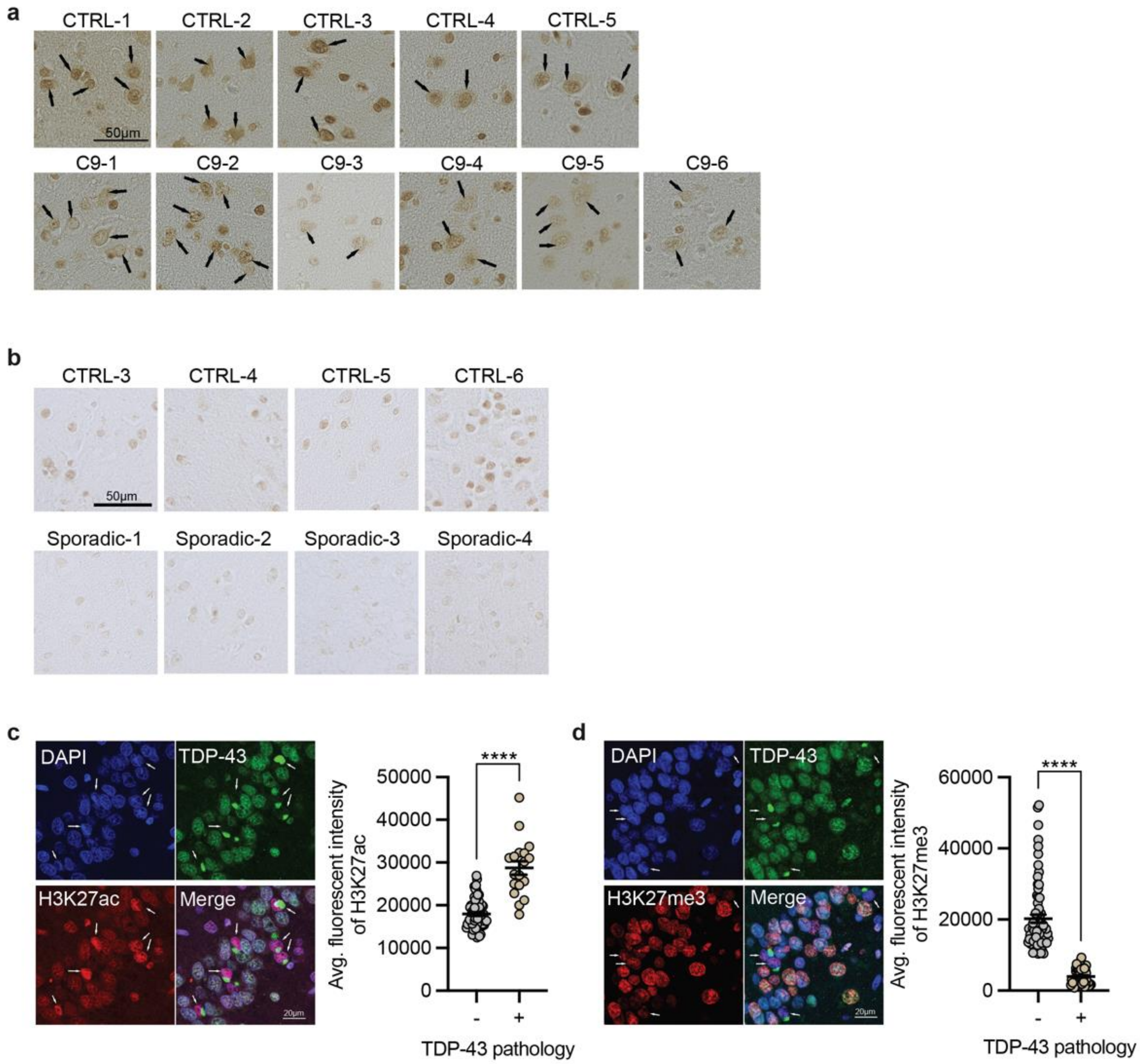


**Extended Data Fig. 5: Chromatin state alteration in patient post-mortem brains.**

**a,** Immunohistochemistry (IHC) staining of H3K9me3 at middle frontal gyrus section of human control and C9ORF72-ALS/FTD samples. Scale bar = 50µm. Arrows point the neurons identified by size and morphology. **b,** IHC staining of H3K9me3 at temporal lobe region of human control and sporadic FTD samples. Scale bar = 50µm. **c-d,** Co-IF labeling and quantification of (c) TDP-43 and H3K27ac, and (d) TDP-43 and H3K27me3 within hippocampus section of a sporadic FTD patient. Arrows point the neurons with TDP-43 pathologies. Scale bar = 20µm. P-values were calculated by two-tailed *t* test on the mean intensities of individual nuclei. Data are mean ± s.e.m. ****p < 0.0001, two-tailed *t* test.

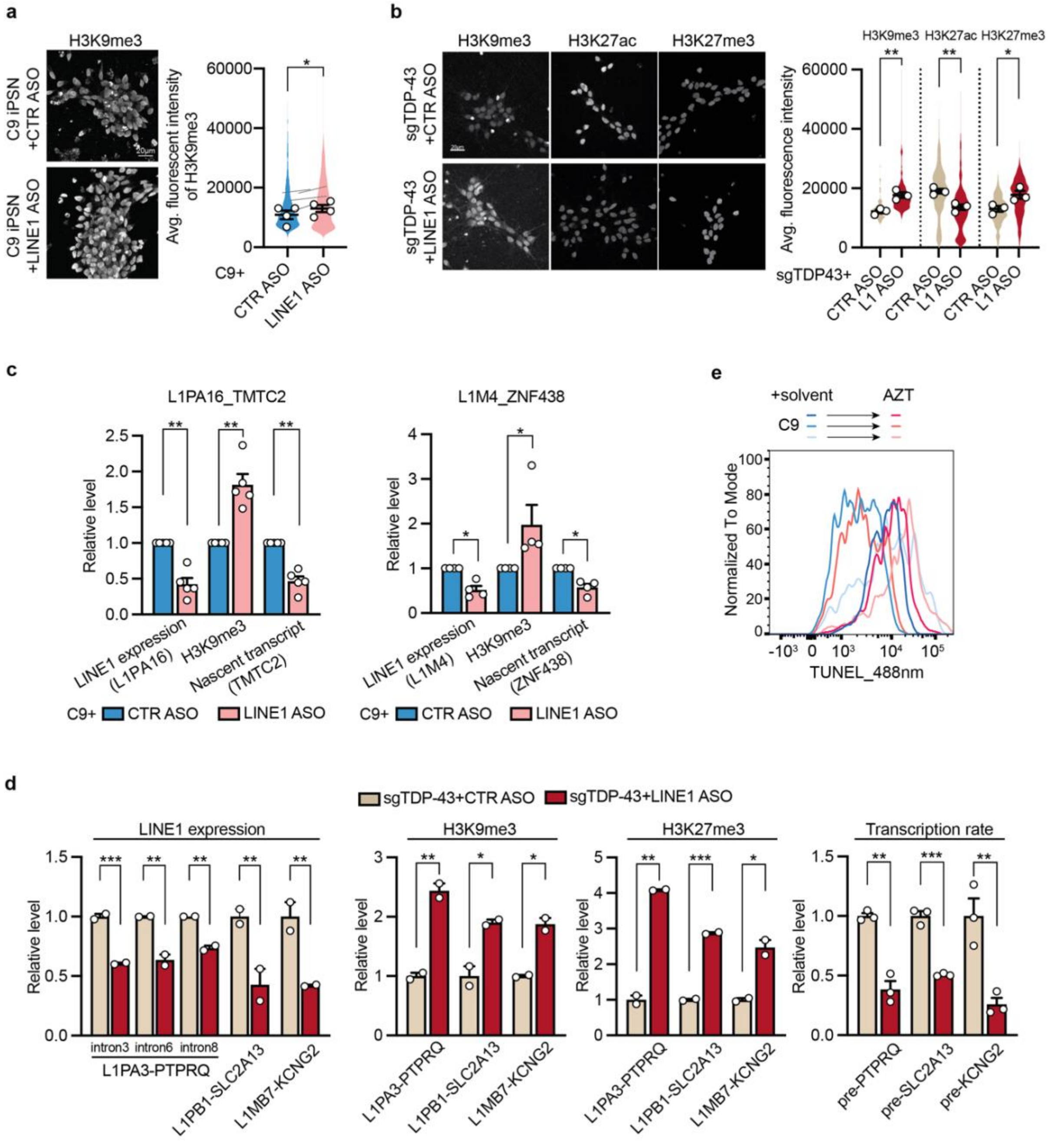


**Extended Data Fig. 6: LINE1 ASO rescues chromatin and transcription abnormalities.**

**a,** IF staining (left) and quantification (right) of H3K9me3 expression in C9 iPSNs treated with control ASO or LINE1 ASO. Points represent individual patient lines with indicated treatment. P-values were calculated by paired two-tailed Student's *t* test on the mean intensities of individual lines. **b**, IF staining and quantification of H3K9me3, H3K27ac, and H3K27me3 levels in TDP-43-reduced neurons following LINE1 ASO treatment. Points represent three biological replicates with indicated treatment. P-values were calculated by two-tailed Student's *t* test on the mean intensities of replicates. **c,** Site-specific LINE1 RNA expression, adjacent H3K9me3 level, and nascent RNA synthesis following LINE1 ASO knockdown in C9ORF72-ALS/FTD iPSNs were examined by RT-qPCR, H3K9me3 ChIP-qPCR, and nascent RNA capture followed by RT-qPCR. Points represent individual patient lines with indicated treatment. P-values were calculated by paired two-tailed Student's *t* test. **d,** Site-specific LINE1 RNA expression, adjacent histone modification levels, and nascent RNA synthesis following LINE1 ASO knockdown in TDP-43-reduced neurons were examined by RT-qPCR, histone-ChIP-qPCR, and nascent RNA capture-RT-qPCR. Points represent individual biological replicates with indicated treatment. P-values were calculated by one-way ANOVA with Tukey's multiple comparisons. **e,** Analysis of chromatin accessibility in patient iPSN lines (n=3 per group) treated with reverse transcriptase inhibitor AZT by DNase I-treated TUNEL assay. Data are mean ± s.e.m. *p < 0.05, **p < 0.01, ***p < 0.001, two-tailed *t* test or one-way ANOVA.

**Supplementary Table 1: Demographic information for iPSC lines.**

| | iPSC line name | Clinical Diagnosis | Age at sampling | Gender | Source |
|---|---|---|---|---|---|
| Ctrl-1 | EDi037-A | Non-neurologic control | 79 | Male | Cedars-Sinai iPSC Core |
| Ctrl-2 | EDi036-A | Non-neurologic control | 79 | Female | Cedars-Sinai iPSC Core |
| Ctrl-3 | CS6EFYiCTR | Non-neurologic control | 50 | Female | Cedars-Sinai iPSC Core |
| Ctrl-4 | CS2AE8iCTR | Non-neurologic control | 50 | Female | Cedars-Sinai iPSC Core |
| Ctrl-5 | CS8PAAiCTR | Non-neurologic control | 58 | Female | Cedars-Sinai iPSC Core |
| Ctrl-6 | 59-1 | Isogenic correction of OXC9-02 | 62 | Female | Kevin Talbot (*1*) |
| C9-1 | CS7VCZiALS | C9orf72 repeat expansion | 64 | Male | Cedars-Sinai iPSC Core |
| C9-2 | CS2GCTiALS | C9orf72 repeat expansion | 64 | Female | Cedars-Sinai iPSC Core |
| C9-3 | CS0NKCiALS | C9orf72 repeat expansion | 52 | Female | Cedars-Sinai iPSC Core |
| C9-4 | C89YHNiALS | C9orf72 repeat expansion | 65 | Female | Cedars-Sinai iPSC Core |
| C9-5 | CS0BUUiALS | C9orf72 repeat expansion | 63 | Female | Cedars-Sinai iPSC Core |
| C9-6 | OXC9-02 | C9orf72 repeat expansion | 62 | Female | Kevin Talbot (*1*) |

**Supplementary Table 2: Demographic information for formalin-fixed paraffin embedded (FFPE) tissue slides from C9ORF72-ALS/FTD and sporadic FTD patients.**

| Tissue ID | Patient GUID | Clinical Diagnosis | Brain region | Age at time of collection | Gender |
|---|---|---|---|---|---|
| Control-1 | BRC2103 | Non-neurologic control | MFG | 88 | Male |
| Control-2 | BRC2052 | Non-neurologic control | MFG | 79 | Male |
| Control-3 | BRC2143 | Non-neurologic control | MFG, STMG | 86 | Male |
| Control-4 | BRC2386 | Non-neurologic control | MFG, STMG | 73 | Female |
| Control-5 | BRC2284 | Non-neurologic control | MFG, STMG | 66 | Female |
| Control-6 | BRC2775 | Non-neurologic control | STMG | 88 | Female |
| C9-1 | BRC1899 | C9ORF72-ALS/FTD | MFG | 86 | Female |
| C9-2 | BRC2589 | C9ORF72-ALS/FTD | MFG | 72 | Female |

| | | | | | |
|---|---|---|---|---|---|
| C9-3 | BRC2665 | C9ORF72-FTD | MFG | 62 | Male |
| C9-4 | BRC2696 | C9ORF72-FTD | MFG | 64 | Female |
| C9-5 | BRC2713 | C9ORF72-FTD | MFG | 70 | Male |
| C9-6 | BRC2786 | C9ORF72-FTD | MFG | 70 | Female |
| Sporadic-1 | BRC2675 | FTD-TDP | STMG | 77 | Male |
| Sporadic-2 | BRC2667 | FTD-TDP | STMG, hippo | 74 | Male |
| Sporadic-3 | BRC2621 | FTD-TDP | STMG | 64 | Female |
| Sporadic-4 | BRC2500 | FTD-TDP | STMG | 63 | Female |

MFG: middle frontal gyrus; STMG: superior temporal gyrus; hippo: hippocampus.

**Supplementary Table 3: Demographic information for frozen postmortem motor cortex tissues.**

| Tissue ID | Patient GUID | Clinical Diagnosis | Gene mutation | Age at sampling | Gender |
|---|---|---|---|---|---|
| Control-1 | JHU100 | Non-neurologic control | NA | 52 | Female |
| Control-2 | JHU101 | Non-neurologic control | NA | 70 | Female |
| Control-3 | JHU108 | Non-neurologic control | NA | 72 | Male |
| Control-4 | JHU123 | Non-neurologic control | NA | 52 | Male |
| Control-5 | JHU124 | Non-neurologic control | NA | 85 | Male |
| Control-6 | JHU129 | Non-neurologic control | NA | 63 | Female |
| C9-1 | JHU22 | fALS | C9ORF72 repeat expansion | 66 | Male |
| C9-2 | JHU88 | fALS/FTD | C9ORF72 repeat expansion | 59 | Male |
| C9-3 | JHU92 | fALS | C9ORF72 repeat expansion | 72 | Male |
| C9-4 | JHU19 | fALS | C9ORF72 repeat expansion | 52 | Male |
| C9-5 | JHU119 | fALS/FTD | C9ORF72 repeat expansion | 61 | Female |
| C9-6 | JHU120 | fALS | C9ORF72 repeat expansion | 68 | Female |

**Supplementary Table 4: Sequences of qPCR primers and ASOs.**

**Supplementary Table 5: High-throughput sequencing sample information.**
